# Optimizing Oscillators for Specific Tasks Predicts Preferred Biochemical Implementations

**DOI:** 10.1101/2022.04.25.489380

**Authors:** Chaitra Agrahar, Michael J Rust

## Abstract

Oscillatory processes are used throughout cell biology to control time-varying physiology including the cell cycle, circadian rhythms, and developmental patterning. It has long been understood that free-running oscillations require feedback loops where the activity of one component depends on the concentration of another. Oscillator motifs have been classified by the positive or negative net logic of these loops. However, each feedback loop can be implemented by regulation of either the production step or the removal step. These possibilities are not equivalent because of the underlying structure of biochemical kinetics. By computationally searching over these possibilities, we find that certain molecular implementations are much more likely to produce stable oscillations. These preferred molecular implementations are found in many natural systems, but not typically in artificial oscillators, suggesting a design principle for future synthetic biology. Finally, we develop an approach to oscillator function across different reaction networks by evaluating the biosynthetic cost needed to achieve a given phase coherence. This analysis predicts that phase drift is most efficiently suppressed by delayed negative feedback loop architectures that operate without positive feedback.

**PACS numbers:** 47.15.-x

## I. INTRODUCTION

While thea literal description of any particular biological system necessarily contains a wealth of molecular detail, a major goal of systems biology is to explain recurring patterns across similar systems in terms of minimal mathematical models. This approach has the advantage of guiding us towards unifying principles between seemingly disparate biological systems. For example, circadian oscillators, in plant [1–5], animals [6–9], fungi [10, 11], and bacteria [12, 13] use quite different molecular components, but share the property that they produce biochemical outputs in a periodic fashion [14]. Basic results from dynamical systems theory indicate that for these systems to produce free-running oscillations they must contain either a multi-step negative feedback loop or a positive feedback loop linked with a negative feedback loop [15, 16]. One approach to simplifying and classifying networks is therefore to refer to them by the number and net logic of feedback loops present. This follows the familiar language of genetic pathway analysis where interactions are identified as positive or negative; these simplified networks are sometimes called topological circuit diagrams [17–19].

However, each abstracted positive or negative interaction in such a diagram represents an underlying biophysical mechanism that impacts the rate of production or removal of the regulated molecule. Concretely, one molecule can negatively influence another by either repressing its synthesis (e.g. acting as a transcriptional repressor) or by stimulating its degradation (e.g. post-translationally modifying a protein to promote its proteolysis). While many possible molecular mechanisms can be envisioned for biochemical oscillators, certain interactions may be found more frequently in natural systems. From the perspective of an evolving system undergoing random mutation, certain mechanisms may be preferred because they are more likely to produce a desired outcome for randomly chosen parameters, and perhaps more easily found by evolution. Here, we pursue a strategy to compare all possible combinatorial regulatory implementations of a given topological diagram (Fig. 1). We use a Monte Carlo approach to sample the molecular constants over a biophysically plausible range of to values to estimate the relative probability of obtaining stable oscillations in the deterministic limit. Later, we introduce an objective function, phase coherence achieved per biosynthetic cost, that allows us to compare the efficiency of suppressing stochastic phase drift across different systems.

**FIG. 1:**
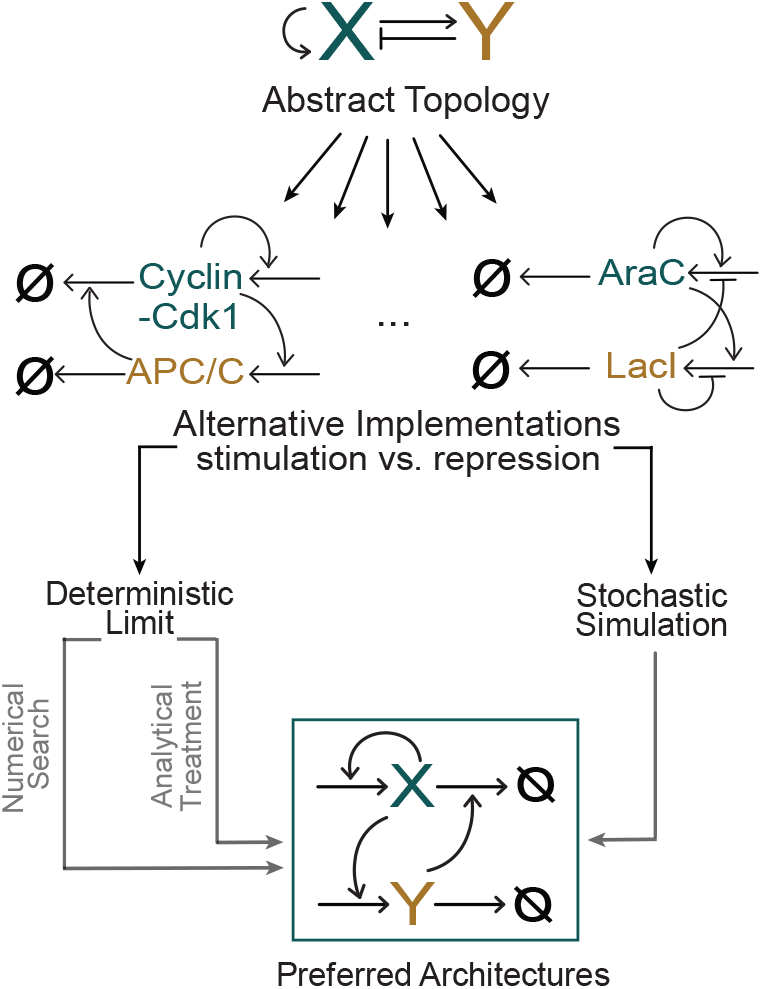
Alternative regulatory implementations of positive and negative interactions are inequivalent in genetic oscillator motifs. We start from the traditional abstract topological description of a positive-negative oscillator motif, and consider all possible regulatory implementations of it. Biological oscillators like the cell cycle with Cyclin-Cdk1 and APC/C as the core components [20] use a different regulatory implementation than synthetic oscillators, like the Hasty Oscillator, with AraC and LacI as core oscillator components [21]. We then compare the relative probability of obtaining stable oscillations using both numerical search and analytical approaches. We compare across systems by evaluating the biosynthetic cost needed to achieve a given phase coherence in the presence of stochasticity.

Although alternative regulatory mechanisms have a similar qualitative effect on the steady state, they have distinct dynamical effects. Fig. 1 shows that there are preferred architectures which have a much higher likelihood of supporting stable limit cycles compared to others, implying that regulatory implementations are more predictive when the criterion of evaluation is the probability of obtaining sustained oscillations.

The importance of considering the regulatory mechanism that implements a feedback loop follows from the basic structure of mass-action kinetics where the net rate of change is a sum of production and removal. Consider a differential equation describing production and removal of a gene product X at constant rates, represented by two opposing terms:

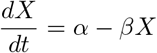

Regulation can be treated in this model by making *α* and *β* non-constant functions of the concentrations of other molecules in the cell, perhaps including X itself. The two terms in this equation are not equivalent for two key reasons. The first is that the basic terms have different kinetic orders in X (one is proportional to X and one is not). The second is that, while a regulated production rate can become arbitrarily small, reflecting tight repression of gene expression, the removal rate constant cannot become smaller than the doubling time of the cell because molecules are diluted by growth.

The general topological description of the systems shown in Fig. 1 is that of a positive-negative oscillator with positive auto-regulation of X, and positive regulation of Y by X, and negative regulation of X by Y. The Hasty oscillator, designed using *E. coli* transcription factors AraC, and LacI, is engineered to have the regulatory implementation as shown in Fig. 1.

The cell-cycle, which is a naturally occurring oscillator has a distinct regulatory implementation compared to the synthetic oscillator, although they share a similar topology. A simplified version of the cell-cycle oscillator is shown in Fig. 1, with two main components: the Cyclin-Cdk1 complex, which stimulates both its own production, and the production of the Anaphase Promoting Complex (APC/C) through Stimulation of Production, hereafter SoP. Cyclin is a cell cycle regulator protein, which binds the Cyclin dependent kinases to activate them and drive the cell cycle forward. The APC/C in turn ubiquitinates the previously active Cyclin-Cdk1 complex leading to degradation of Cyclin B. That is, the negative APC/C arm of the oscillator acts negatively through Stimulation of Degradation, or SoD.

Fig. 1 illustrates the approaches used to determine the regulatory mechanism which maximizes the likelihood of oscillations in the deterministic and the stochastic limit. To statistically analyze the properties of reaction networks within this scheme, we randomly sample parameters according to the following rules:

I. the unregulated production reactions are zeroth order, and the unregulated degradation reactions are first order in the gene product.
II. positive and negative regulation are modeled so that the unregulated rate constants for production or degradation become functions of the concentration of other molecules. We assume that the regulatory interactions are monophasic, so the modifying functions are monotonic.
III. There are unregulated basal production (*α*_0_) and degradation (*β*_0_) rates. Modifying functions act on non-basal production (*α*) and degradation (*β*) rates.
IV. the modifying functions which act to enhance the rate constants in the numerical simulations have a Hill function form, given by 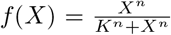. The comparing oscillatory circuits of different architecture by determining the protein synthesis cost required to achieve a given phase coherence. modifying function which suppresses the rate constants have the form 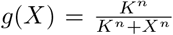, where *K* is the affinity parameter, and *n* is the Hill coefficient, such that *n >* 0 indicates co-operativity between molecules.
V. when two regulatory functions act on the same step, they are treated additively, following the treatment described in [22].
VI. the regulation of production can be tight, i.e. expression of a gene may be completely dependent on a given transcription factor, meaning basal production rates can go to zero. However, basal degradation rates can never be zero due to the dilution factor introduced by the growth of cells. So, basal degradation rate *β*_0_ *> k*_*growth*_.
VII. both production and degradation rates have upper bounds reflecting the finite time taken for the production or degradation of a macromolecule, i.e.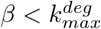 and 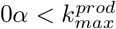.

The results are organized as follows: We consider all possible combinatorial regulatory implementations of a topological diagram in the deterministic limit. By Monte Carlo sampling the molecular constants underlying a given regulatory mechanism, we quantitatively evaluate system function for a given objective function. We mainly focus on two 2-node systems, and one 3-node system (topological diagrams with 2 or 3 interacting genes or gene products), though the approach can be easily generalized to larger systems.

We then present a study of the loop oscillator and the positive-negative oscillator motifs in the stochastic limit. We show that the phase coherence in the stochastic limit can be estimated through the deterministic arclength in the phase-space of the oscillating components. We outline the role of symmetry in determining the upper bound for phase-coherence of oscillations at a given deterministic orbit arclength. We develop a metric for comparing oscillatory circuits of different architecture by determining the protein synthesis cost required to achieve a given phase coherence.

Finally, we discuss relevant examples from biological systems and demonstrate the application of our predictions. We also consider the differences in the design of biological oscillators and synthetic oscillators to understand how to design synthetic oscillators with higher phase coherence of oscillations. We extend our analysis of loop oscillators to networks of post-transcriptional loop oscillators in Appendix B3.

## II. RESULTS

### Positive-Negative Feedback Oscillators

Many oscillators contain interlocking positive and negative feedback loops. That is, components of the oscillator act in two ways: they promote their own activity through one pathway while ultimately leading to inhibition of their activity through another pathway. Many oscillators contain interlocking positive and negative feedback loops. That is, a component of the oscillator act in two ways: it promotes its own activity through one pathway while ultimately leading to its inhibition through another pathway. A minimal version of this system consists of two components: a self-activating positive element, and a negative element which participates in a negative feedback loop (Fig. 1). We implement the positive regulation in two distinct ways: i) Stimulation of the Production reaction (SoP), Repression of the Degradation reaction (RoD). The negative regulation is implemented in two distinct ways: through the Stimulation of Degradation (SoD), or through the Repression of Production (RoP).

Applying the principles listed in the Introduction, we show that this result can be understood intuitively in terms of a phase plane analysis. When the positive element acts much faster than the negative element, conditions for oscillation can be seen graphically using phase plane analysis. Below we work in this separation of timescales limit—we will conclude that regions of the phase plane become comparatively inaccessible for certain regulatory schemes.

We first analyze the self-regulation of X and the positive regulation of Y by X. To predict the implementations which maximize the probability of obtaining stable limit cycles, we ultimately combine the constraints on the each individual regulatory step. For the positive element (X), a necessary condition for oscillation is that this nullcline has local extrema, and the nullcline for the negative element (Y) must cross in the switchback region between them (Fig. 2A). This nullcline structure implies that there is a range of Y values where, if Y were held constant, the self-activation of X must give bistable dynamics.

**FIG. 2:**
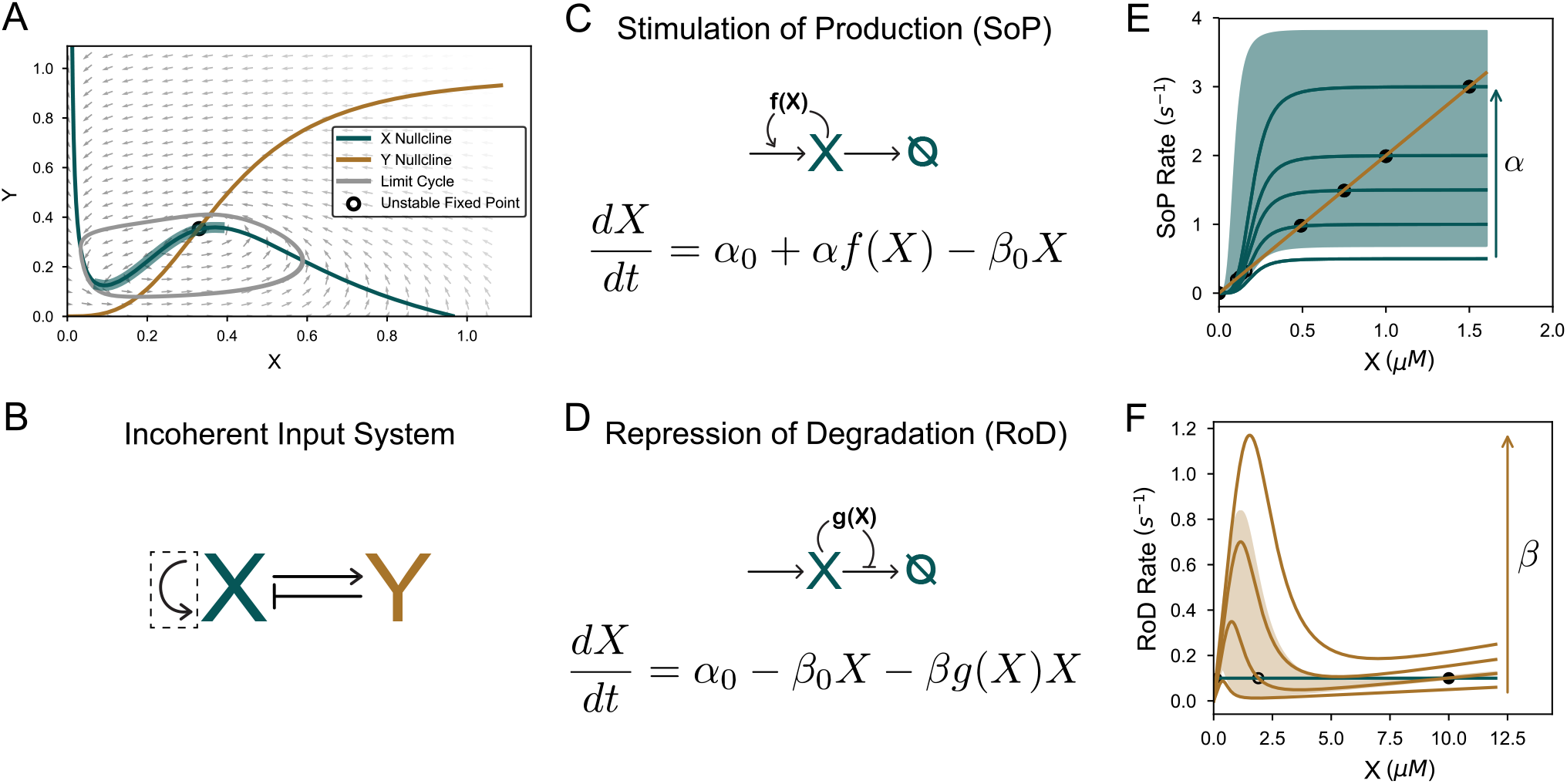
Stimulation of Production (SoP) is the preferred implementation for the positive self-loop. *α*_0_, and *β*_0_ are the basal unregulated production and degradation rates. A) Illustrates the X and Y nullclines. To obtain stable limit cycles, the X and Y nullclines must cross in the switchback region of the X nullcline (thick teal part of the X Nullcline). The grey curve indicates the limit cycle, and the arrows denote the vector field flow. B) The highlighted positive autoregulatory interaction of X (dashed box) is implemented through SoP. C) The positive autoregulation of X is implemented through SoP. D) The positive autoregulation of X is implemented through RoD. E) Region of bistability supported when positive autoregulation of X is implemented through SoP is large. When *α*_0_ is negligible, a stable fixed point near zero exists for any value of *α*. Similarly,The fixed point at zero is always stable, as is the fixed point at high values of X is stable (filled circles). However, the intermediate fixed point is unstable (open circles). The bistability of SoP is robust to changes in production (teal line) and degradation rates (yellow line). F) The region of bistability supported when positive autoregulation of X is implemented through RoD is comparitively small.

### Positive self-regulation by X

As bistable dynamics of X, for fixed Y are required to obtain stable limit cycles, we start by asking which implementation of self-activation is more likely to give a large region of bistability, for X (Fig. 2B). The SoP implementation where a monotonically increasing function *f* (*X*) regulates the production rate *α* (Fig. 2C), is logically compared to the RoD implementation where a monotonically decreasing function *g*(*X*) regulates the production rate *β* (Fig. 2D). The stimulatory function *f* (*X*) can go to zero, meaning regulation is tight. However, there is a maximum production rate 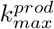, which bounds *f* (*X*) above. The repressive function *g*(*X*) cannot be lower than the minimum degradation rate *β*_0_, which is the dilution factor due to cell growth *k*_*growth*_. The maximum degradation rate *β*, bounds *g*(*X*) above.

By varying the production and degradation rates, and the parameters of the regulatory functions through random sampling, we conclude that SoP implementation of the positive self-regulation by X leads to a larger region of bistability on average (Figs. 2E and F). In figs. 2E and 2F, the fixed point at small values of X is always stable, as is the fixed point at high values of X (filled circles). However, the intermediate fixed point is unstable (open circles). The bistable dynamics of SoP exists though a larger range of variations in the production (teal line) and degradation rates (yellow line), and the parameters of the regulatory functions. The bistability of RoD is conditional on the value of the Hill co-efficient such that 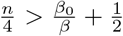, and tolerates a much smaller range of variations in production and degradation rates, and other system parameters.

### Analysis of X Nullclines

A Mote Carlo search through the parameter space described in Table 1, for all possible regulatory implementations of a topological diagram reveals that numerically one implementation vastly outperforms every other implementation in terms of maximizing the probability of supporting stable limit cycles (Fig. 1, and see below). There is one preferred implementation which maximizes the probability of obtaining stable limit cycles. A mechanistic understanding of this result can be obtained through a geometrical analysis of how nullclines interact in the phase space.

**TABLE I:** Experimentally motivated distributions of parameter from which the function parameters are drawn [17], [23]-[24].

| System Parameter | Ranges Used |
| --- | --- |
| $\alpha_0, \beta_0$ | $[10^{-3}, 10^{-2}] \mu\text{M min}^{-1}, \text{min}^{-1}$ |
| $\alpha, \beta$ | $[10^{-3}, 10^2] \mu\text{M min}^{-1}, \text{min}^{-1}$ |
| $K$ | $[10^{-3}, 10^2] \mu\text{M}$ |
| $n$ | $[1, 4]$ |

In the limit where the X dynamics are faster than the Y dynamics, i.e.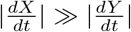, a necessary condition for oscillation is that the X nullcline has local extrema, and the Y nullcline crosses only in the switchback region, described in Fig. 3A. To predict which regulatory implementation maximizes the probability of a nullcline crossing only in the switchback region, we study the geometry of both the X and Y nullclines.

**FIG. 3:**
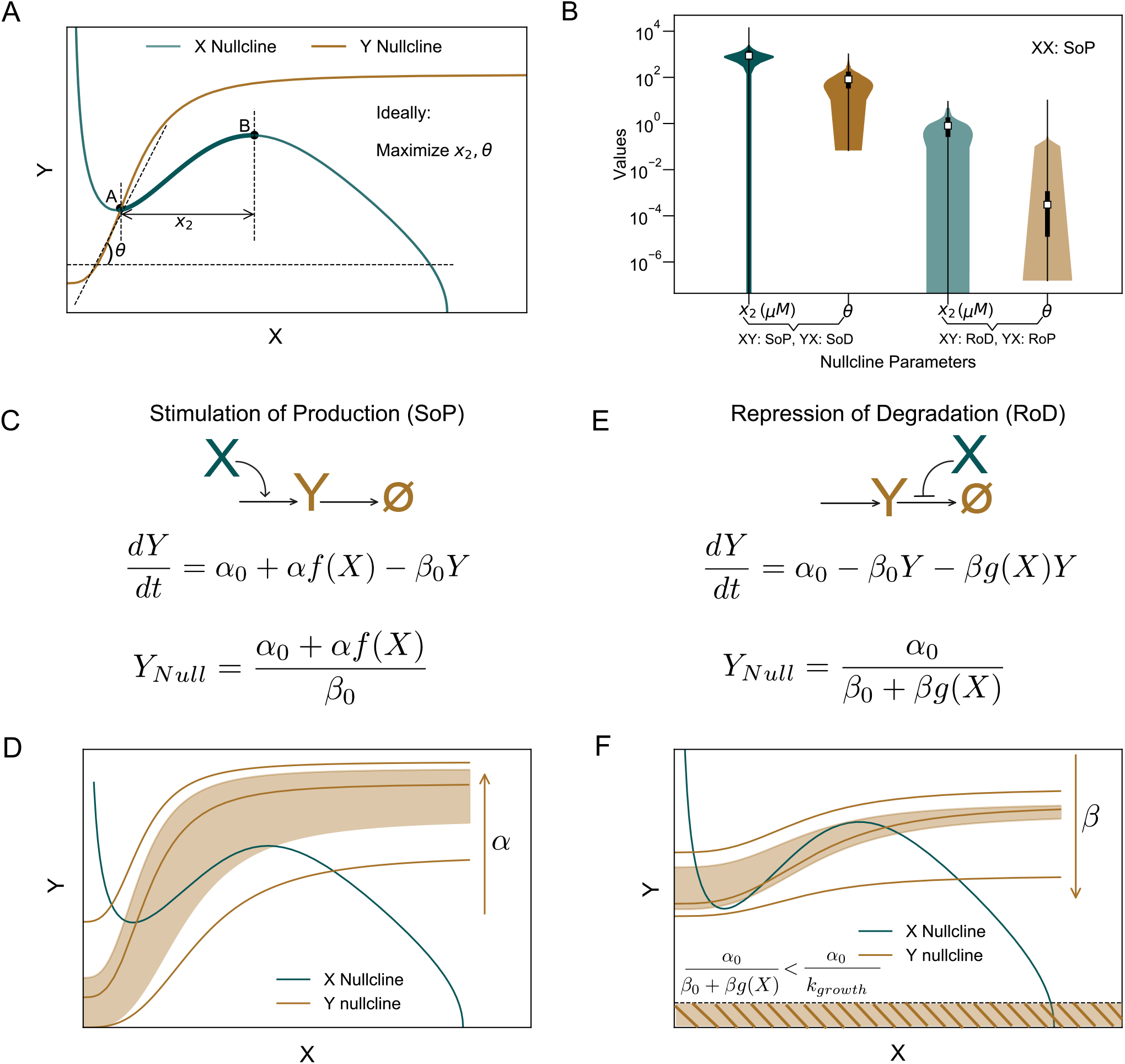
SoP is the optimal implementation of the positive regulation if Y by X. A) Limit cycles are obtained when the Y nullcline crosses the X nullcline in the switchback region (between points A and B). The probability of obtaining limit cycles can be estimated through geometrical parametrization of the nullclines. To maximize the likelihood of obtaining limit cycles, the switchback region of the X nullcline, *x*_2_, and the slope of the linearized region of the Y nullcline, *θ* both should be maximized, along with other geometrical constraints are described in the Supplement (SI.9). B) shows the distribution of the *x*_2_ and *θ* values for 10,000 random parameter sets, each implemented in the two different ways described in C) and E). C) and E) show the analytical expressions for the Y nullclines, as implemented through SoP and RoD, respectively. Shaded regions of the graphs indicate the possible Y nullclines, which can produce stable limit cycles in D) and F). Stimulation of Production (SoP) is the preferred implementation for the positive regulation of Y, compared to the Repression of Degradation (RoD) implementation, as illustrated by the shaded regions in D) and F).

The optimal geometry for X nullclines can be determined through four parameters. The most predictive of the four parameters, is the width of the switchback region of the X nullcline, *x*_2_, shown in Fig. 3A. Maximizing *x*_2_ increases the region available for the crossing of the Y nullcline within the switchback region, thereby increasing the probability of obtaining stable limit cycles.

### Analysis of Y Nullclines

The geometry of the Y nullcline can also be characterized by four geometrical parameters. The most predictive of these is the slope of the linearized region of the Y nullcline *θ*, shown in Fig. 3A. To obtain stable limit cycles, *θ* should not be too small, or the Y nullcline will be below the X Nullcline, and cross it outside the switchback region, after the second local extremum. On the other hand, *θ* cannot be too large, as it will lead the Y nullcline to be too steep, and cross the X-nullcline to the left of the switchback region, before the first local extremum.

Thus, the regulatory implementation which maximizes the likelihood of the Y nullcline crossing the X Nullcline within the switchback region is one which puts *θ* in the range described in Equation Box 1.

### Crossing of X and Y Nullclines

In addition to having in the right range of values, it is necessary to have constraints on the geometry of the Y nullclines such that the crossing with the X nullcline occurs in the switchback region, and not outside of it. This includes constraints on the minimum and maximum points at which the Y nullcline can begin to rise, and where it plateaus. *θ*, and three other parameters characterize the geometry of the Y nullcline. The geometrical constraints on these parameters, and their combined effects on the geometry of the Y nullcline, and the crossing between the X and Y nullclines are discussed in Appendices B-E, and in SI 2. We show numerically, and through analytical means that the SoP implementation of the positive regulation of Y by X increases the probability of a stable limit cycles. Numerically, as can be seen in Fig. 3B, median *x*_2_ values in the SoP scheme can be 3 orders of magnitude higher than the median *x*_2_ values for the RoD implementation of the interaction.

Analytically, as shown in Fig. 3C, we see that the Y nullcline is proportional to the regulatory function describing X stimulating production of Y. By the principle that regulation can be tight, this structure implies that the Y nullcline can access the biochemical zero, which implies a tight regulation of the production rate. Fig. 3D illustrates cases where the Y nullcline can access the biochemical zero. This feature of the regulatory implementation enables enables the nullclines to enter a wider region of the phase-space, and increases the likelihood of obtaining stable limit cycles.

In contrast, we can see in Fig. 3E, that the repressive regulatory function acting on the degradation rate is in the denominator of the Y nullcline. For a growing cell, this establishes a lower bound of 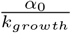 for the region of phase-space accessible to the Y nullclines in the RoD implementation. Fig. 3F illustrates the region of phase-space that is inaccessible due to the lower bound established by the growth rate of cells. As the Y nullclines are constrained to a value above 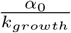, for the X and Y nullclines to cross in the switchback region, the slope of the linearized region must be smaller on average, compared to the SoP implementation. However, this also implies that the region of crossing becomes smaller, as shown in Fig. 3F, and leading to a smaller probability of obtaining stable limit cycles.

In addition to increasing the width of the switchback region of the X nullcline, it is also necessary to have constraints on the geometry of the nullclines such that the crossing of the X and Y nullclines occurs only in the switchback region. The geometrical constraints on these parameters, and their combined effects on the geometry of the X Nullcline, and the crossing between the X and Y nullclines are discussed in Supplementary Figure SI. 2. The principles laid out in the Introduction help us predict the optimal implementation for two node systems analytically (Appendix Supplementary Methods).

**Equation Box 1:**
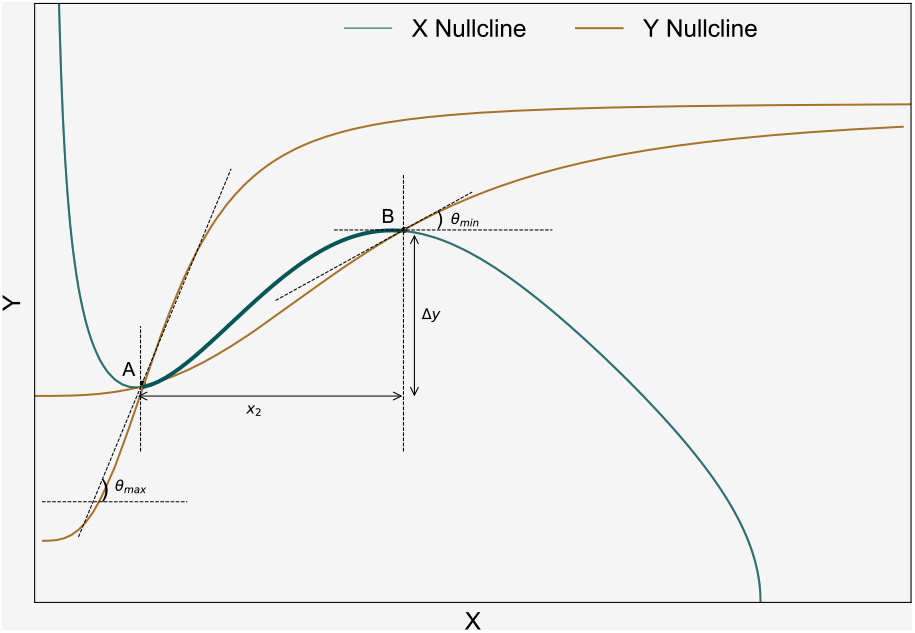
Ranges of slopes of linearized Y nullclines that allow for stable limit cycles.

The range of slopes for the Y nullclines using Table 1.

**SoP:**

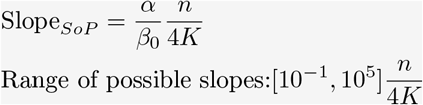

**RoD:** If *n* = 4,

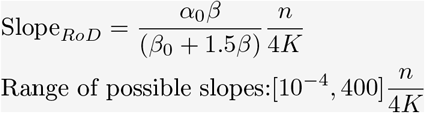

For details of the calculations of slope, see Methods M6.

Stable limit cycles occur when the Y nullclines cross the X nullcline in the switchback region, denoted by *x*_2_. However, merely increasing the switchback region does not necessarily increase the probability of obtaining stable limit cycles if the geometry of the nullclines are incommensurate with a crossing in the switchback region. This is because the geometrical constraints on both the shape of the nullclines, and the geometrical constraints on the crossing which determine the probabilities rather than just one of the two factors. To obtain stable limit cycles, the Y nullclines must cross within the rectangle determined by *x*_2_ and Δ*y*. This implies that the linearized slope of the Y nullclines must fall in the range of values between *θ*_*min*_ and *θ*_*max*_.

### Dual Regulation is the optimal implementation of a single-input negative interaction

The negative loop of the positive-negative oscillator motif can be structured in two different ways: one where the node X receives two inputs with contradictory logic (Incoherent Input system, inset Fig. 4A), and one where the node X receives two inputs with similar logic (Coherent Input system, inset Fig. 4B) [17].

**FIG. 4:**
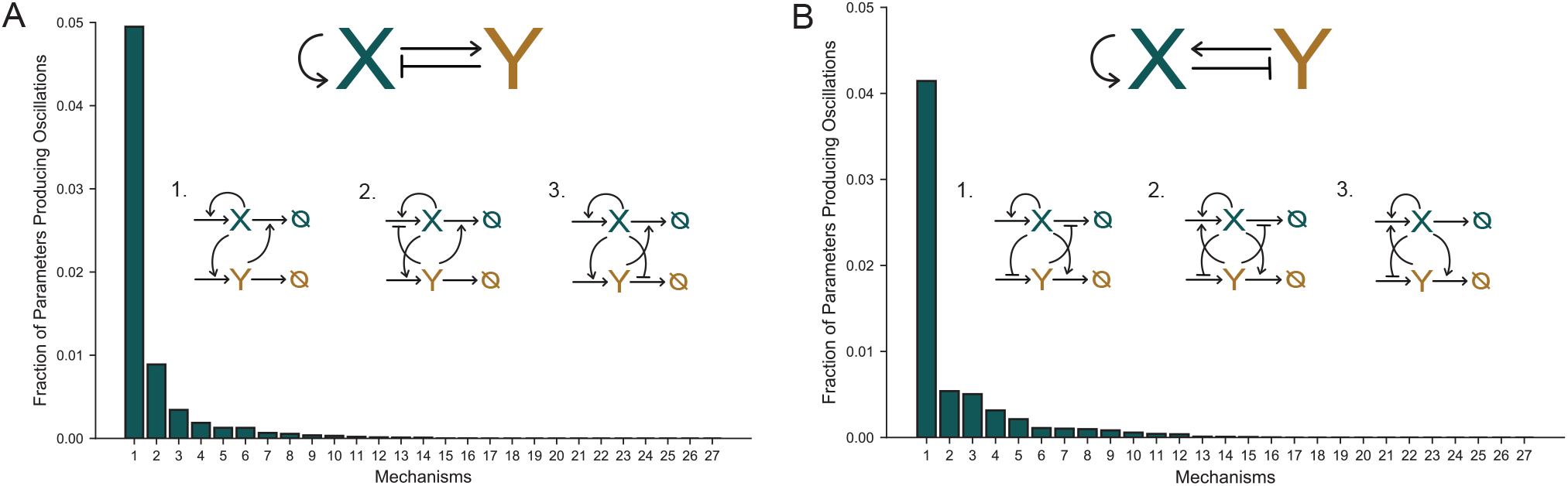
The likelihood of robust oscillations depends strongly on the regulatory implementation. A) Incoherent input system topology (top inset). The bar chart quantifies the robustness responses of each of the 27 distinct implementations of the incoherent input topology. Each implementation results in a different robustness response of the oscillator topology, of which the top three most robust implementations are shown here. B) Coherent input system topology (top inset). The bar chart quantifies the robustness responses of each of the 27 distinct implementations of the coherent input topology. Each implementation results in a different robustness response of the oscillator topology, of which the top three most robust implementations are shown here. We compare the bar charts in A) and B) to conclude that coherent and incoherent input systems have comparable robustness when each is implemented with the optimal mechanism, as shown in the insets in the above panels.

Up to this point we have considered mechanisms where a molecule only regulates one of the reactions. We now consider the possibility that both reaction steps may be regulated, which we term “dual regulation”. We treat each arm of the dual regulation independently, meaning, the affinity parameters for each step of the regulation are chosen independently from the distribution in Table 1. Combinatorically, each interaction of the topological diagram can be implemented molecularly in three different ways, leading to 27 distinct regulatory implementations. Numerically simulating all possible implementations of the positive-negative oscillator motif, we find that one implementation has a much higher probability of supporting stable limit cycles compared to all others (Figs. 4A-B).

For incoherent input systems described in Fig. 4A, the preferred implementation is through stimulation reactions. Namely, the positive interactions are implemented as the stimulation of production, and the negative interaction is implemented as the stimulation of degradation. For coherent input systems to have a similar likelihood of obtaining stable limit cycles, the positive interactions are implemented through stimulation of production. However, the negative interaction implemented as a dual regulation, acting on both production and degradation steps maximizes the probability of sustained oscillations.

Previous work has shown that incoherent input systems enhance the likelihood of obtaining stable oscillations, compared to coherent input systems [17]. However, these analyses all made the apriori choice to implement all interactions through stimulation mechanisms. We see here that allowing repression mechanisms in the negative feedback arm drastically increases the probability of oscillation for the coherent topology. In particular, if the negative interaction in the coherent input system is implemented through dual regulation (Fig. 4B), the probability of obtaining stable limit cycles is comparable to the best implementation of the incoherent input system (Fig. 4A).

### Loop Oscillators

While positive-negative motifs are one way to achieve oscillations, it is also possible to create oscillating biochemical circuits with multistep negative feedback loops that do not utilize positive feedback. A historically important model of a biological oscillator is the Goodwin model, a negative feedback-only loop, where the high degree of cooperativity in the final negative interaction provided the necessary non-linearity to achieve oscillations [25]. The repressilator motif (Fig. 5A (i)), created synthetically by Elowitz and Leibler [26] distributes the non-linearity over multiple steps, and can thus generate oscillations with a smaller degree of cooperativity in any individual step. In the repressilator loop, each component is a transcription factor that inhibits expression of the next element in the loop. Thus, negative logical interactions are achieved through repression of production (RoP).

**FIG. 5:**
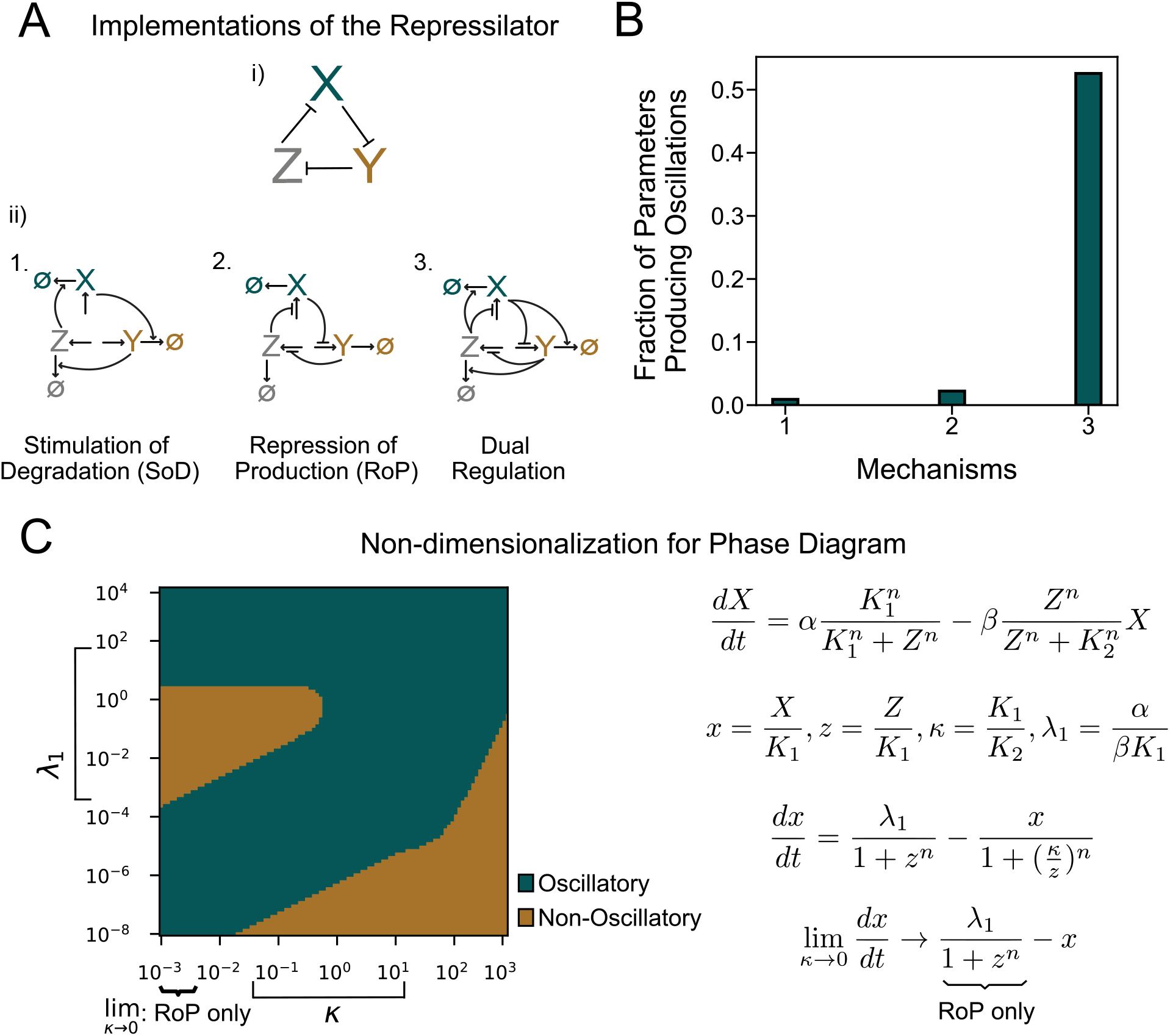
Dual regulation is the optimal implementation of negative regulation when a node receives only one negative input. A) The repressilator motif (i) is implemented in three different ways, through (ii) 1. the Stimulation of Degradation (SoD), 2. the Repression of Production (RoP), and 3. Dual Regulation. B) The bar chart shows the fraction of parameter sets that give stable limit cycles when parameters are drawn randomly from distributions of the reaction rates of the repressilator motif. C) Phase diagram, with *n* = 3, depicts the region of stability of a dual regulated repressilator, non-dimensionalized such that the *κ →* 0 limit recovers the regulation through repression of production limit. Square brackets in panels indicate the biologically plausible region for a growing *E. coli* cell, from which the parameter sets are drawn to arrive at the bar chart in panel B). The mathematical implementation of the non-dimensionalization which gives the phase diagram in C) is also presented.

Following the same methodology used above to scan over regulatory interactions, we consider a simplified version of the repressilator, without explicitly modeling the transcription process. A simplified mathematical description of one of the genes of the repressilator circuit can be written as:

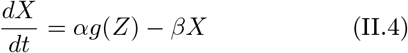

where *g* is the repressive regulatory function defined in the Introduction. However, we can also model the negative interaction as Z stimulating the degradation rates of In this case the dynamics of X can be written as:

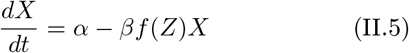

where *f* is the stimulatory regulatory function, described in the Introduction.

Each of the two implementations of the repressilator discussed above achieve negative regulation by acting on different reaction steps. The negative feedback can be implemented through the repression of the production step (RoP), or through the stimulation of the degradation step (SoD), or through Dual Regulation. The three implementations of the Repressilator motif are shown in Figs. 5A ii (1-3).

For each implementation of the loop shown in Fig. 5A, we quantify the fraction of parameter sets randomly drawn from the distributions in Table 1 which give stable limit cycle oscillations. Fig. 5B compares the numerical outcomes in each case. Notably, while RoP is preferred compared to SoD, Dual Regulation, where each component can regulate both production and removal is markedly superior. Implementation of the negative interaction through dual regulation is five times more likely to give oscillations than other implementations of the negative interaction. This result is commensurate with our finding in Fig. 4B, which shows that in coherent input systems, at the node Y which receives a single negative input, Dual Regulation enhances the probability of obtaining stable limit cycle oscillations.

We confirm numerically and analytically, that Dual Regulation enhancing the likelihood of obtaining stable limit cycles is not an artefact of a specific choice of parameter distribution or the particular form of the Hill function used to model regulation. Analytically, for symmetric systems, where the rate constants associated with the three dynamical variables are identical, we derive the condition for oscillations through the Jacobian, and verify that the condition for oscillations for the Dual regulated system is more likely to be satisfied, as it constructively combines the conditions for each of the two single regulations. This is described in more detail in Equation Box 2. Numerically, we consider two non-dimensionalization schemes, which have the property that we obtain one of the two single regulation limits as we tune a system parameter *κ* (Fig. 5C) to zero. One such scheme, where we recover the single regulation mechanism where the negative regulation is achieved through the repression of the production step, is shown in Fig. 5C.

**Equation Box 2:**
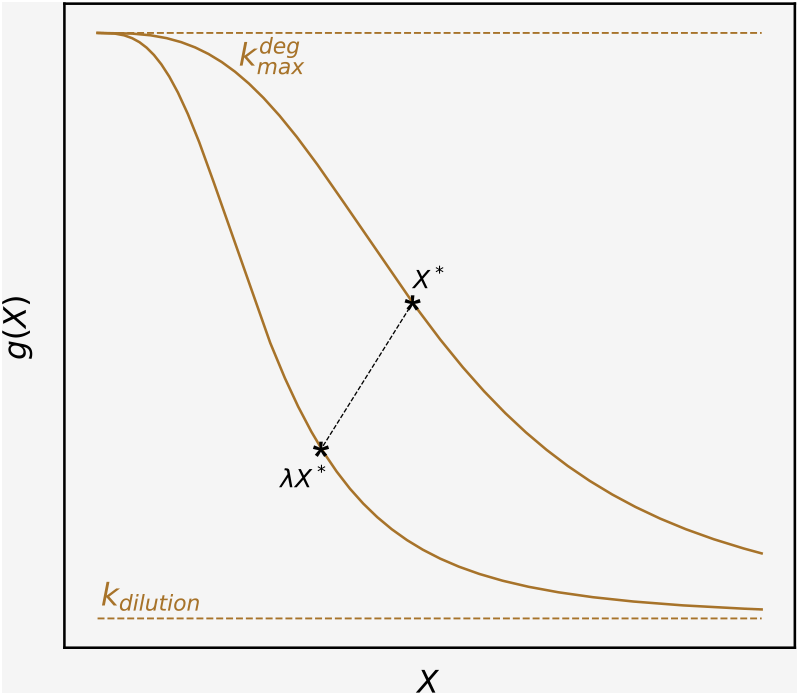
We compare the conditions for oscillations for the three implementations of the repressilator system. Linear stability analysis gives us the condition for oscillations when the negative loop is implemented through the repression of the production reaction:

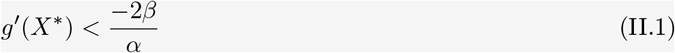

where *g*^′^(*X*^∗^) is the slope of the repressive regulatory function *g* at the fixed point *X*^∗^, *α* is the production rate, and *β* is the degradation rate.

The condition for oscillations when the repressilator loop is implemented through the stimulation of the degradation reaction is:

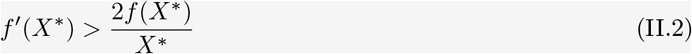

where *f* ^′^(*X*^∗^) is the slope of the stimulatory regulatory function *f* at the fixed point *X*^∗^.

The reason that RoP is more likely to give oscillations can be understood through a scaling argument. If the fixed point *X* is rescaled by a scalar *λ* such that *X*^∗^ → *λX*^∗^, then *g*^′^(*λX*^∗^) → *λg*^′^(*X*^∗^), which rescales the above equation. So, if one set of parameters do not satisfy the condition initially, a rescaling factor *λ* can be found such that Eqn. 1.1 is satisfied, assuming the position of the fixed point is set by the scale of *g*(*X*).

However, such a rescaling factor scales both sides of Eqn. 1.2. Thus, the probability of obtaining stable limit cycles is higher for RoP implementation compared to the SoD implementation.

The condition for oscillations for dual regulation of the repressilator is:

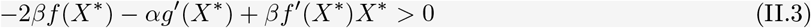

For most parameter sets, we can find a rescaling factor *λ* such that parameters satisfying (1.1) or (1.2) also satisfy Eq (1.3), making dual regulation the easiest way to obtain oscillations.

The phase-diagram showing the regions of parameter space that support oscillations is shown in Fig. 5C. For each randomly drawn parameter set, we calculate the Jacobian, and verify whether that parameter set satisfies the condition for oscillation or not.

We see that the regions of phase-diagram that support oscillations expand as we increase *κ*. As we approach the Dual Regulation regime, the probability of obtaining stable limit cycles increases. Dual Regulation also performs progressively better as we increase the value of the parameter *κ*, in the case where *κ* → 0 limit recovers the single regulation where negative interaction is implemented through Stimulation of Degradation (see SI 6).

### Regulatory implementations in biological and synthetic oscillators

To contextualize our numerical and analytical results for positive-negative oscillators, and loop oscillators, we consider multiple biological and synthetic oscillators which have been characterized experimentally. We sketch out the regulatory mechanisms of the interactions that have been discovered experimentally. Biological oscillators that may be used to generate precise timing likely require a high degree of phase-coherence and tend to utilize the incoherent input system topology, and its preferred implementation, XX: SoP, XY: SoP, and YX: SoD.

Synthetic oscillator systems are typically built such that the regulations are all on the production step. This is presumably because it is experimentally easier to engineer circuits by introducing transcription factor binding sites to stimulate or repress production reactions using a cognate transcription factor in the circuit. However, our results show that implementing the negative regulation through the stimulation of the degradation will lead to a higher probability of sustained oscillations in an incoherent input system.

In the following discussion, we present simplified architectures for some natural biological oscillators, and suggest regulatory implementations for enhancing the operational range of synthetic oscillators. For natural systems where the molecular architecture appears to deviate from the optimal designs described here, our results may motivate a search for previously undescribed regulatory interactions. This is particularly the case for dual regulation (both production and removal are regulated) which has the potential to markedly enhance oscillator function and may have been missed in previous experimental studies.

Many biological oscillators like the mammalian circadian oscillator, the cell cycle, etc. when represented in their simplest forms, share the incoherent input positive-negative system topology [2], [27]. Here we observe that the implementation of this regulatory logic is also shared across many natural systems. Specifically, a positive element that stimulates its own production, and a negative element that stimulates the removal or degradation of the positive element.

### The mammalian circadian oscillator

The mammalian circadian clock is complex, and involves multiple components that make it tick. We consider a simple model based on previous work, where the mammalian circadian oscillator is modelled as being composed of four genes, or their protein products, namely Clock, Bmal1, Cry and Per [27]. The circadian oscillator functions through two complexes of heterodimers formed out of these four proteins: the BMAL1-CLK complex, and the PER-CRY complex.

The mammalian circadian oscillator consists of positive and negative feedback arms which function together to give stable 24 hour oscillations. BMAL1 and CLOCK proteins form a heterodimer, which then binds to the E-box promoters of PER and CRY genes, and activate their transcription [27], through SoP.

Per and Cry proteins, in turn form a heterodimer complex, and translocate into the nucleus. An increased concentration of CRY in the nucleus results in a binding of CRY to the BMAL1-CLK-E-box complex, inhibiting the activity of BMAL1-CLK complex [27] - [30] through SoD. Notably, the PER/CRY complex does not direct act to repress transcription. Rather, PER/CRY converts BMAL1-CLK into an inactive form. Although this reaction does not destroy BMAL1-CLK, the rate of transforming active BMAL1-CLK into an inactive complex depends on CRY, making the kinetic equations equivalent to SoD.

Finally, an increased concentration of PER in the nucleus dissociates the BMAL1-CLK complex, and induces the transcription of BMAL1 by repressing the NRd1 factors, which inhibit BMAL1 transcription [31], [30], and positively autoregulating through SoP.

The regulatory implementations utilized in the mammalian oscillator is the most robust implementation of the positive-negative incoherent input system. Similarly, the cell cycle, and the pulsatile actomyosin contractility in *C. elegans* all implement the interactions in their shared topology through the implementation that maximizes the likelihood of obtaining stable limit cycles for the incoherent input system topology, as shown in Fig. 1A [20], [27], [29], [32].

### Synthetic Oscillators

Synthetic oscillators have been built for various purposes – repressilator [26], ComK-MecA (SynEx) oscillator [29], Hasty Oscillator [21] etc. The common element among the synthetic oscillators is that the regulatory implementations of the logic is achieved by altering the gene expression levels using transcriptional promoters. Positive regulations are implemented via the stimulation of gene expression, and negative regulations are implemented via the repression of gene expression.

While oscillations can be achieved this way, our results indicate that the probability of obtaining stable limit cycles will increase in an incoherent input architecture when positive regulations are implemented through stimulation of production steps, and negative interactions are implemented by targeting the degradation machinery.

An example demonstrating this is readily found in [29], where the authors re-engineered the ComK-ComS system bypassing the pathway of ComS inhibiting the degradation of ComK by MecA. The resulting “SynEx” circuit has incoherent input logic, and is implemented through the preferred regulatory mechanisms identified here. Experimentally, this change in the circuit from a non-preferred implementation of the coherent input system to the preferred implementation of the incoherent input system results in its loss of variability compared to the native competence circuit with ComK and ComS.

This leads us to hypothesize that the biological function influences both the choice of topology and regulatory implementation, as the native ComK - ComS circuit likely benefits from variability, functioning to produce population heterogeneity in response to stress [29]. To further demonstrate this hypothesis, we now introduce stochasticity into the analysis.

### Oscillator Motifs in the Stochastic Limit

So far, we have studied oscillator dynamics in the deterministic limit, and we have evaluated circuit function based on the probability of obtaining stable oscillations. This corresponds to the biological limit of having infinitely many copy numbers of the proteins that form the biochemical circuits. If Ω denotes the system size, where we imagine scaling up the size of the cell and gene copy number together, then lim Ω → ∞ is the deterministic limit where all effects can be described in terms of continuous concentrations. When Ω is finite, stochastic effects become noticeable. Cells have finite volume and thus biochemical systems in the cell are composed of finitely many proteins. This may be especially important for bacteria where a protein present at a concentration of 1 nM corresponds to approximately one protein copy in a typical *E. coli* cell. An alternative design criterion for oscillators is therefore their resistance to stochastic noise. Here we develop an approach to compare noise resistance across oscillator architectures.

We study the different regulatory implementations of the repressilator motif and the positive-negative oscillator motif in the stochastic limit. We use the Gillespie stochastic algorithm to compute the time series of the stochastic oscillator dynamics. Fig. 6A shows deterministic (bold line) and stochastic trajectories (thin lines) for the incoherent input system (Fig. 6A (i)), and the RoP Repressilator system (Fig. 6A (ii)). In the stochastic limit, one measure of performance for an oscillator is the phase-coherence of stochastic oscillations. We define phase-coherence as the height of the first non-unity peak of the autocorrelation function of the stochastic time series. Fig. 6B shows the deterministic and stochastic phase-space orbits. The procedure used to calculate the phase-coherence of oscillations is described in Supplement Methods M2.

**FIG. 6:**
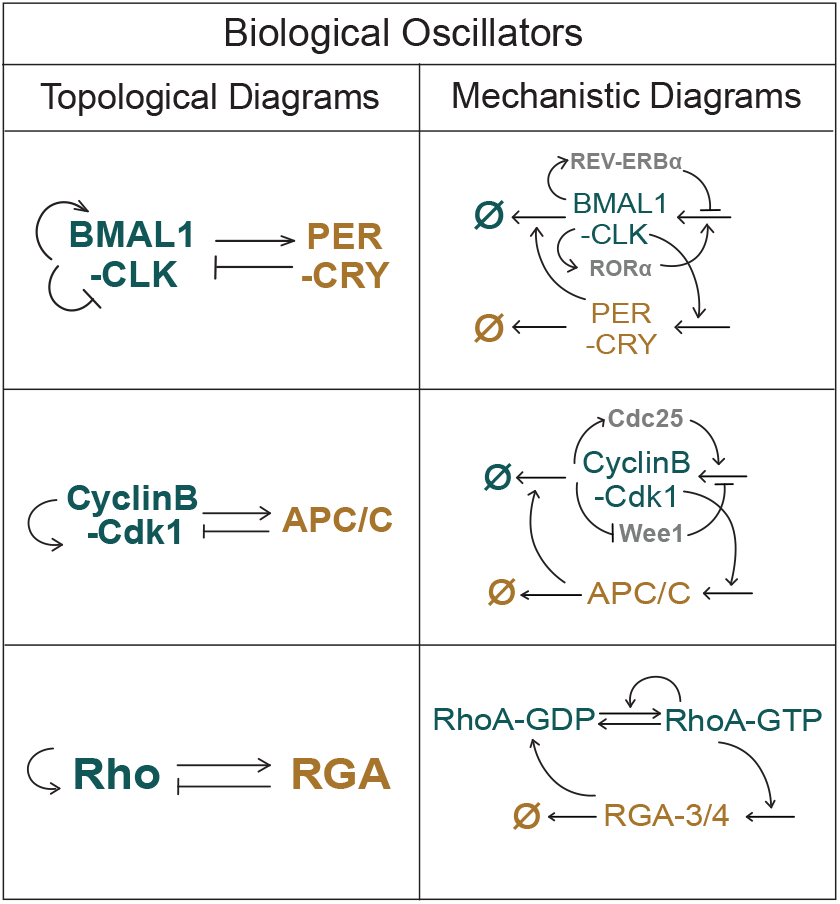
Table 2a: Models of biological oscillators in literature for the mammalian clock [27], for the cell cycle oscillator18, and for the rho-rga system in *C. Elegans* [28] are depicted in Column I. Column II illustrates the mechanisms through which the interactions occur.

**FIG. 7:**
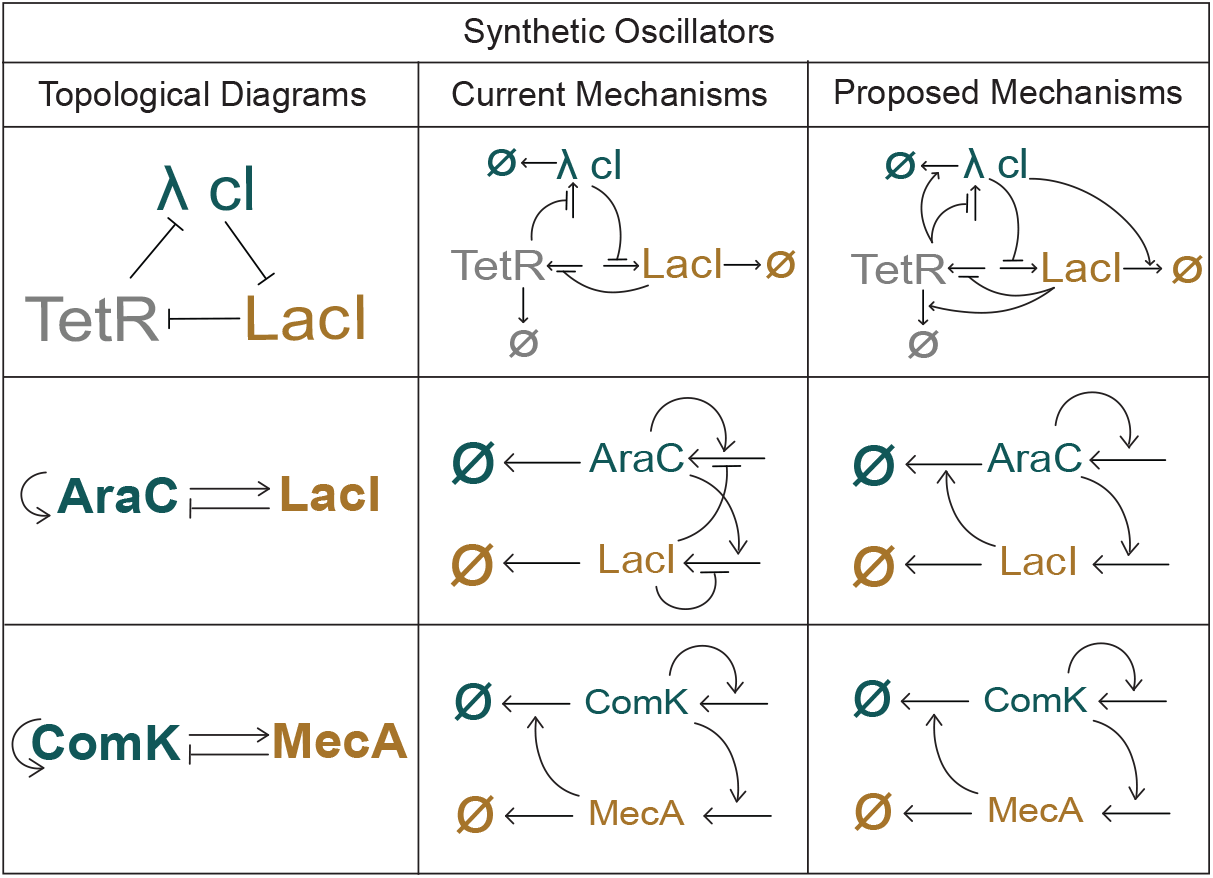
Table 2b: Examples of synthetic oscillator systems. Column I shows the topologies of some synthetic oscillator systems. Column II depicts the regulatory mechanisms as they are currently implemented experimentally [26],[21],[29]. Column III illustrates the proposed optimal regulatory implementations for the topologies in Column II.

**FIG. 8:**
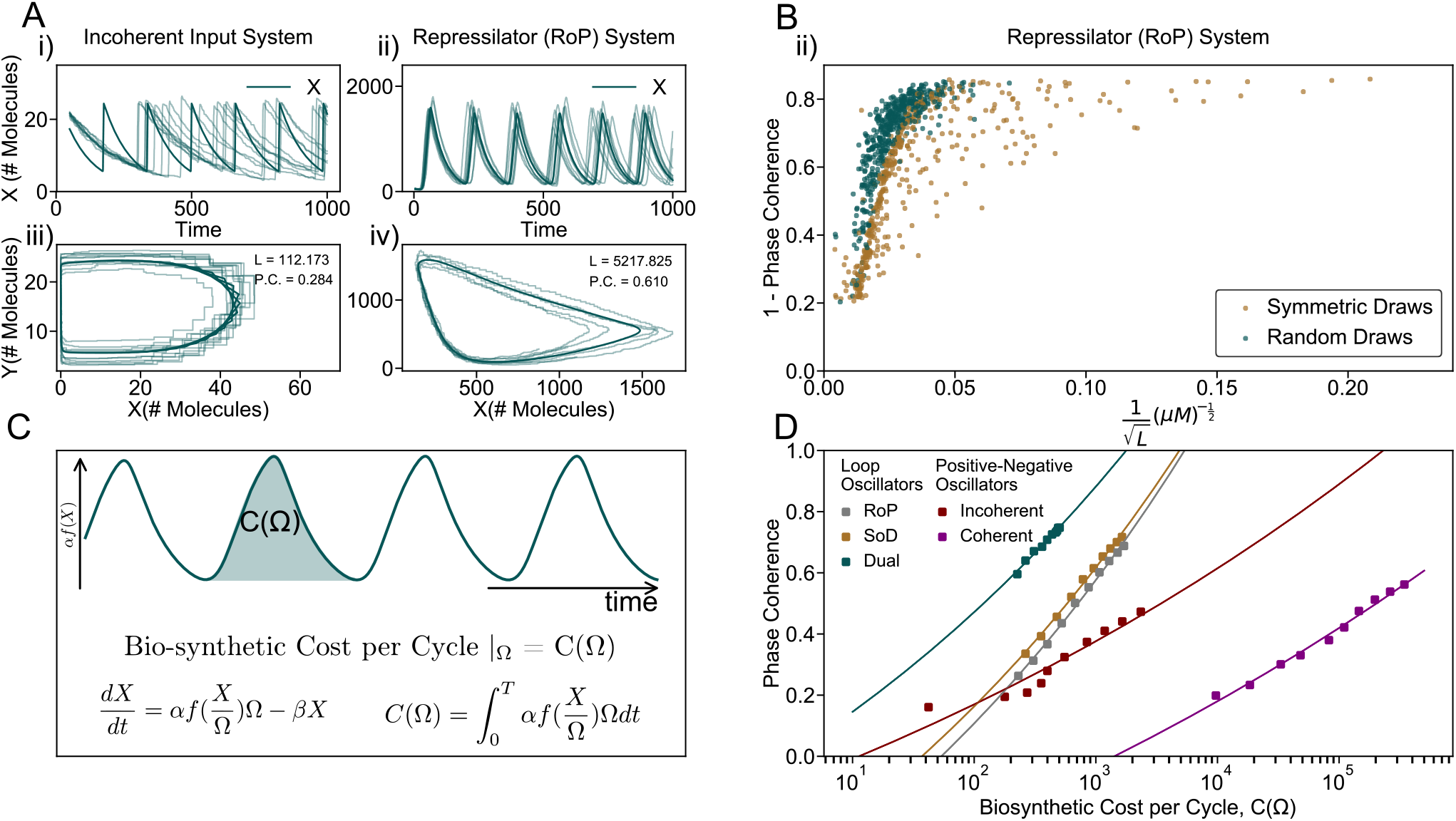
Loop oscillators optimize the trade-off between biosynthetic cost and phase-coherence of oscillations. A) Deterministic (bold lines) and stochastic (thin lines) simulations of the most robust implementation of the i) incoherent input positive-negative oscillator time series, and ii) the repressilator motif with negative interaction implemented through RoP, in a symmetric system time series. Phase space orbits of iii) the incoherent input system, and iv) the repressilator. B) Relationship between the deterministic arclength (L) of the phase-space orbit, and the stochastic phase coherence for the i) incoherent input system, and ii) the repressilator. The vertical axis represents the error in phase incurred in one oscillation cycle. Symmetric parameter sets (teal dots), (i.e. repressilators with the same rate parameters for all dynamical variables) saturate the upper bound for phase coherence values at a given deterministic arclength and protein copy number. Asymmetric parameter sets (yellow dots) generally do not saturate the phase-coherence-deterministic arclength relationship. To compare the performance of different topologies and regulatory implementations of those topologies, we define the parameter-free metric of the biosynthetic cost per cycle needed to achieve a certain phase coherence value of oscillations. C) Illustrates the notion of biosynthetic cost per cycle of oscillation, at a given copy number,. This is the integral of the production rate over the period of oscillation. D) The biosynthetic cost per cycle needed to achieve different values of phase-coherence for the three loop oscillator implementations, and the most robust implementations of the coherent and incoherent input system topologies. The square dots represent the cost per cycle for different values of, as obtained through an iterative process towards reaching a value of phase coherence equal to 0.7. Each square marker represents the median value of phase-coherence and biosynthetic cost for 130 sets of parameters. We see that the loop oscillator systems achieve higher phase coherence values at lower costs when compared to positive-negative oscillator systems.

We consider the repressilator motif, which has a symmetric topology, with cyclic repression of the genes in the loop. The fully symmetric version of this system has the same biochemical parameters for all the genes in the loop.. We also consider asymmetric parameter sets within the repressilator motif, which retain a symmetric topology, but have asymmetric interaction strengths. Both the symmetric and the asymmetric realizations of the repressilator are implemented in the three ways discussed in the section on loop oscillators.

Fig. 6B shows a relationship between the length of the deterministic phase-space orbit, and the error incurred over one cycle of oscillation. This is consistent with the intuition that oscillations with a larger deterministic orbit (higher amplitude) will comprise more elementary reaction steps in the stochastic limit; fluctuations in the time to complete the orbit can be suppressed by averaging over more steps.

Numerically, we observe that there is a lower bound on the error incurred over a cycle of oscillation, which is set by symmetric systems. For systems and parameters where velocity around the orbit is constant, this relationship between the deterministic arclength and the phase-coherence value can be a good estimate for the stochastic behavior of the system. Thus, negative feedback-only loop oscillators perform optimally when timescale and interaction strength is matched across all the components

### Biosynthetic Expenditure - Phase-Coherence Trade-off

Our analysis of different regulatory implementations relied on a comparison of the relative probability of obtaining stable oscillations for random draws of parameters. It is difficult to generalize this approach to make comparisons between different classes of biochemical systems that may have very different numbers of parameters or draw those parameters from distinct distributions.

Here we propose a method to compare oscillator function in the stochastic limit across architectures that circumvents these problems. We suppose that the important function of the circuit is to produce regular oscillations (i.e. to minimize phase error) and that it operates subject to a constraint of finite resources. Thus for each oscillator architecture we can define the maximal possible phase coherence for a given biosynthetic investment per cycle. Although all oscillators achieve perfect regularity with infinite resources, the scaling with finite resources can be quite different (Fig. 6). Fig. 6D shows the relationship between the biosynthetic cost and the phase-coherence value that can be achieved for a given topology and implementationA clear result is that the negative feedback-only loops are more efficient at achieving high phase coherence. The dual regulation scheme, where each element in the loop acts on both production and degradation of the next, is markedly superior to either alone.

Considering the positive-negative feedback oscillator systems, we see that the optimal implementation of the coherent input system requires a much larger biosynthetic expenditure to achieve a given phase-coherence value, compared to the optimal implementation of the incoherent input system. Although these two systems can have similar probabilities of giving deterministic oscillations from random parameter sets, the coherent scheme gives markedly noisier oscillations.

This gives a rationale for why biological oscillators where high phase-coherence is likely desirable—like the mammalian circadian oscillator—would utilize incoherent connections between positive and negative feedback loops, while oscillators where high phase-coherence is not the desired functionality, like the stochastic state switching ComK-ComS circuit in *B. subtilus* utilizes the coherent input system [21].

## III. DISCUSSION

Biological oscillators like the cell cycle, the circadian sleep-wake cycle, etc. are integral to the survival of organisms. They regulate downstream processes and confer a fitness advantage by anticipating the time of day, even in the absence of external cues [28], [33]. Thus, decoding the design principles of oscillator circuits is crucial to understand the workings of biological oscillator circuits, as well as for designing robust synthetic oscillators.

Tikhonov and Bialek point out in their paper that the focus on network topology as the sole determiner of biological function is, in all likelihood a result of the developmental arc of experimental genetics [34]. It is harder to study the mechanistic pathways involved in the execution of a biological function than it is to infer the net logic of interaction between two known participant genes. This experimental bottleneck has resulted in the study of biological function as the product of circuit topologies rather than of the pathways of implementation of the logic encoded in topological diagrams.

Although many circuits can sustain robust oscillations, few network motifs have been evolutionarily conserved. We see the recurrence of loop oscillator motifs and motifs of positive-negative oscillators in many systems, and on many different timescales of oscillations [17]. The functional advantage of these motifs is attributed to the enhancement of the robustness of oscillations [17], [22], [35]. The ability to reliably function over a wide range of parameter values is a natural consideration, as organisms routinely need to function in variable environments.

example, Tsai et. al. show that the addition of positive self-loops to negative feedback cores enhances the probability of obtaining oscillations [22]. Positive feedback loops also allow for high amplitude oscillations over a wide range of frequencies [22], [35]. After systematic analyses of all possible two and three node circuit topologies, Li et. al. arrive at the conclusion that joining inputs incoherently at a node, where a positive and a negative input type feed into the node lead to enhanced probability of oscillations, as compared to coherent input systems where two positive or two negative inputs feed into a single node [17]. These studies analyze models within a fixed paradigm where all positive and negative interactions are implemented through stimulation of a reaction channel.

We verify their results, but find that the function of a given topology depends on crucially on the mechanism by which the logical interactions are implemented. By searching over all possible implementations, we find that positive-negative systems with a coherent logic can have similarly robust function to incoherent logic systems provided they are implemented with an optimal mechanism. This is fundamentally because different implementations lead to different structures of the kinetic equations. The preferred molecular mechanisms can therefore be predicted by studying the structure of the ODEs in phase space, and through linear stability analysis.

Remarkably, the performance and robustness of negative feedback loop systems is dramatically improved when the possibility of dual regulation is allowed so that each element acts on both the production and removal step of the next element. In concrete terms, dual regulation could be implemented by a transcription factor that represses expression of its target, while also inducing expression of a protease that specifically degrades the target. Whether biology uses dual mechanisms that have so far remained hidden in natural oscillators is an interesting open question.

To compare performance of different circuits, potentially with very different structure, we introduce a parameter-free measure. We compare the biosynthetic cost for maintaining oscillations with a given value of phase coherence. As cells are generally resource limited, this measure allows us to find the implementation that minimizes the biosynthetic expenditure for maintaining high phase coherence of oscillations. While interlocked positive and negative feedback loops are known to have a wide operational range in terms of parameters that support oscillation, we find that pure negative feedback loop oscillators are superior when judged by resistance to stochasticity. Again, the highest performing system is the negative feedback loop implemented through dual regulatory mechanisms. While real biological systems are more complex than the small circuits studied here, it is straightforward to extend this phase coherence metric.

Based on these results, we predict that different implementations of a topology are preferred for different biological functions. In general, biological oscillators may be under selection for functions that are quite distinct. One possible function is time-keeping, where frequency and phase stability are key to maintain correlation with an external signal, e.g. in a circadian rhythm. Another possible function is to ensure that the system reliably progesses through a sequence of distinct states with high amplitude and that the process recurs—possibly a description of the free-running cell cycle. Here, a small amount of phase drift may be tolerable. A third possible function is for an oscillator to intentionally operate in the stochastic regime for the purpose of creating heterogeneity in a population. Our results suggest that these distinct functions will be mirrored by alternative underlying molecular architectures For systems that require e.g. a pulse generator where high phase coherence is unnecessary (or even undesirable), positive-negative oscillators will be preferred. However, for applications such as biological clocks where a stable phase relationship between the oscillator and the external environment is key, negative feedback loops may be preferred.

Finally, our results suggest that synthetic biology efforts have often used sub-optimal circuit designs to create oscillators. Future work engineering targeted protein or RNA degradation may open up the ability to control the degradation terms in the kinetic equations, likely enhancing the robustness of oscillator function. The identification of topological oscillator motifs has been a huge step forward for systems biology, but our understanding of the molecular mechanisms typically used to implement these motifs remains incomplete. Our work provides a mechanistic understanding of the robustness of different biological oscillator motifs, finding that the choice of which reaction steps are regulated is a crucial determinant of function. The approach developed here can be extended to study how the regulatory implementations in driven oscillator systems can alter the robustness versus entrainment trade-off in oscillator systems. The end goal of this line of analysis will be to determine whether specific oscillator mechanisms are preferred depending on the functional demands and fitness constraints on the system.

## Supporting information

Supplemental Infomation

## IV. ACKNOWLEDGEMENTS

CA thanks the Physics Department at UIC for the teaching stipend that enabled this work. We thank members of the Rust lab, Dr. Moumita Das, and Dr. Martin Falk for their inputs on the manuscript. This work was supported by an HHMI Faculty Scholar award and NIH Grant R01 GM107369 to MJR.

## Data Availability

Code used for the numerical analysis can be found at chaitraagrahar.github.io. This study includes no data deposited in external repositories.

## Author contributions

Chaitra Agrahar: Conceptualization; Simulation; Formal analysis; Visualization; Methodology; Writing—original draft; Writing—review and editing. Michael Rust: Conceptualization; Formal analysis; Supervision, Writing—review and editing. The contributions in detail are: CA and MJR conceived and designed the study. CA developed the computational pipelines and analyzed data under supervision of MJR. CA wrote the manuscript.

