## Supplemental Infomation for "Optimizing Oscillators for Specific Tasks Predicts Preferred Biochemical Implementations"

(Dated: April 25, 2022)

PACS numbers: 47.15.-x

**Contents**


**APPENDIX** 3

**A. Condition for Oscillations in the Repressilator** 3

1. Condition for oscillations in the Repression of Production (RoP) Mechanism 3
2. Condition for oscillations in the Stimulation of Degradation (SoD) Mechanism 4
3. Condition for oscillations in the Dual Regulation Mechanism 5

**B. Generalized analysis for post-translational and transcriptional loop oscillator systems.** 6

1. Condition for oscillations for generalized oscillator systems repression of production (RoP) or repression of activation (RoA) 6
2. Condition for oscillations for generalized oscillator systems stimulation of degradation (SoD) or stimulation of deactivation (SoD) 7
3. Condition for oscillations for generalized oscillator systems with dual regulation 8

**C. Nullcline analysis for incoherent input systems** 8

1. Determining the optimal regulatory implementations when Y is negatively regulated through the stimulation of the degradation rate 9
2. Determining the optimal regulatory implementations when Y is negatively regulated through the repression of the production rate 13

**D. Nullcline analysis for coherent input systems** 15

**E. Nullcline analysis of the mammalian circadian oscillator** 19

**APPENDIX SUPPLEMENTARY METHODS** 21

**M1. Convergence Criteria for limit cycles** 21

**M2. Calculation of Phase Coherence Values** 21

**M3. Positive-Negative Oscillator Motifs** 22

**APPENDIX SUPPLEMENTARY FIGURES** 28

### APPENDIX

#### Appendix A: Condition for Oscillations in the Repressilator

We derive the condition for oscillations through the linear stability analysis of the three-node transcriptional repressilator when the repression is implemented via the mechanism of repression of the production reaction. The repressilator is symmetric in this underlying motif, meaning that there is a cyclic repression executed by the three genes X,Y, and Z. For analytical tractability we also assume that the interaction strengths between the three genes are identical, making the system completely symmetric.

##### 1. Condition for oscillations in the Repression of Production (RoP) Mechanism

A regulatory implementation of the repressilator where repression is implemented through modulation of the production rates can be written as:

$$\begin{aligned} \frac{dX}{dt} &= \alpha g(Z) - \beta X \\ \frac{dY}{dt} &= \alpha g(X) - \beta Y \\ \frac{dZ}{dt} &= \alpha g(Y) - \beta Z \end{aligned} \tag{A.1}$$

where $g$ is a regulatory function that inhibits production rates. A typical example of $g(Z)$ would be

$$g(Z) = \frac{K^n}{(K^n + Z^n)} \tag{A.2}$$

where $K$ is a measure of the binding affinity, and n denotes the degree of cooperativity among $Z$ molecules required for the reaction to occur.

Let us assume that the fixed point of the system is $fp = (X^*, Y^*, Z^*)$. As the parameters are assumed to be symmetric, the three dynamical variables are interchangeable. This, in turn allows the system to have a fixed point that is also symmetric, i.e. $fp = (X^*, X^*, X^*)$. At the fixed point, the Jacobian can be evaluated as:

$$J_{(X^*,X^*,X^*)} = \begin{bmatrix} -\beta & 0 & \alpha g'(X^*) \\ \alpha g'(X^*) & -\beta & 0 \\ 0 & \alpha g'(X^*) & -\beta \end{bmatrix} \tag{A.3}$$

The condition for the existence of an unstable fixed point is that the real part of the eigenvalues of the Jacobian must be positive. If $\lambda$ is the eigenvalue, then the condition for oscillations is that $Re(\lambda) > 0$. The condition for oscillation for a production regulated repression mechanism impose a constraint on the slope of the regulatory function:

$$g'(X^*) < \frac{-2\beta}{\alpha} \tag{A.4}$$

This condition is easy to satisfy, and the likelihood of obtaining oscillations higher. To see why, consider a choice of parameters which doesn't satisfy the condition (A1.4). Then, a rescaling of the regulatory function by some non-zero, non-negative scalar $\lambda$ rescales the LHS of the Eqn (A1.4).

Scaling by $\lambda$, if $X \to \lambda$ X, then the fixed point goes to $X^* \to X^*/\lambda$. By chain rule, $g(X) \to g(\lambda X)$. The derivative scales by $\lambda$, i.e. $g'(\lambda X) \to \lambda g'(X)$. On scaling, the condition for oscillations becomes:

$$\lambda g'(X^*) < \frac{-2\beta}{\alpha} \tag{A.5}$$

A careful choice of the rescaling factor $\lambda$ can help satisfy the condition for oscillations for this system, as rescaling only changes the slope of the regulatory function, and allows for higher likelihood of oscillations.

##### 2. Condition for oscillations in the Stimulation of Degradation (SoD) Mechanism

We derive the condition for oscillations through the linear stability analysis of the three-node transcriptional repressilator when the repression is implemented via the mechanism of stimulation of the degradation reaction.

For analytical tractability we also assume that the interaction strengths between the three genes are identical, making the system completely symmetric. A regulatory implementation of the repressilator where repression is implemented through modulation of degradation rates can be written as:

$$\begin{aligned} \frac{dX}{dt} &= \alpha - \beta f(Z)X \\ \frac{dY}{dt} &= \alpha - \beta f(X)Y \\ \frac{dZ}{dt} &= \alpha - \beta f(Y)Z \end{aligned} \tag{A.6}$$

where $f$ is a regulatory function that inhibits production rates. A typical example of $f(Z)$ would be

$$f(Z) = \frac{Z^n}{(K^n + Z^n)} \tag{A.7}$$

where $K$ is a measure of the binding affinity, and n denotes the degree of cooperativity among $Z$ molecules required for the reaction to occur.
Let us assume that the fixed point of the system is $fp = (X^*, Y^*, Z^*)$. As the parameters are assumed to be symmetric, the three dynamical variables are interchangeable. This, in turn allows the system to have a fixed point that is also symmetric, i.e. $fp = (X^*, X^*, X^*)$. At the fixed point, the Jacobian can be evaluated as:

$$J_{(X^*,X^*,X^*)} = \begin{bmatrix} -\beta f(X^*) & 0 & -\beta f'(X^*)X^* \\ -\beta f'(X^*)X^* & -\beta f(X^*) & 0 \\ 0 & -\beta f'(X^*)X^* & -\beta f(X^*) \end{bmatrix} \tag{A.8}$$

The condition for the existence of an unstable fixed point is that the real part of the eigenvalues of the Jacobian must be positive. If $\lambda$ is the eigenvalue, then the condition for oscillations is that $Re(\lambda) > 0$. The condition for oscillation for a degradation regulated repression mechanism imposes a constrain on the slope of the regulatory function:

$$f'(X^*) > \frac{2f(X^*)}{X^*} \tag{A.9}$$

This condition is not so easy to satisfy, and the likelihood of obtaining oscillations lower. To see why, consider a choice of parameters which doesn't satisfy the condition (A2.4). Then, a rescaling of the regulatory function by some non-zero, non-negative scalar $\lambda$ rescales the LHS of the Eqn (A2.4).

Scaling by $\lambda$, if $X \to \lambda X$, then the fixed point goes to $X^* \to X^*/\lambda$. By chain rule, $f(X) \to f(\lambda X)$. The derivative scales by $\lambda$, i.e. $f'(\lambda X) \to \lambda f'(X)$. On scaling, the condition for oscillations remains:

$$f'(X^*) > \frac{2f(X^*)}{X^*} \tag{A.10}$$

Rescaling the regulatory function doesn't help satisfy the condition for oscillations for this system, as rescaling changes the both sides of Eqn (A2.5). hence, a choice of parameters which initially doesn't satisfy the condition for oscillations on scaling will still not give oscillations.

##### 3. Condition for oscillations in the Dual Regulation Mechanism

We derive the condition for oscillations through the linear stability analysis of the three-node transcriptional repressilator when the repression is implemented via both the repression of the production reaction and the stimulation of the degradation reaction. A regulatory implementation of the repressilator where repression is implemented through dual regulation can be written as:

$$\begin{aligned}\frac{dX}{dt} &= \alpha g(Z) - \beta f(Z)X \\ \frac{dY}{dt} &= \alpha g(X) - \beta f(X)Y \\ \frac{dZ}{dt} &= \alpha g(Y) - \beta f(Y)Z\end{aligned} \tag{A.11}$$

where $f$ and $g$ are regulatory functions used in appendices A2 and A3. that inhibits production rates.

Let us assume that the fixed point of the system is $fp = (X^*, Y^*, Z^*)$. As the parameters are assumed to be symmetric, the three dynamical variables are interchangeable. This, in turn allows the system to have a fixed point that is also symmetric, i.e. $fp = (X^*, X^*, X^*)$. At the fixed point, the Jacobian can be evaluated as:

$$J_{(X^*,X^*,X^*)} = \begin{bmatrix} -\beta f(X^*) & 0 & \alpha g'(X^*) - \beta f'(X^*)X^* \\ \alpha g'(X^*) - \beta f'(X^*)X^* & -\beta f(X^*) & 0 \\ 0 & \alpha g'(X^*) - \beta f'(X^*)X^* & -\beta f(X^*) \end{bmatrix} \tag{A.12}$$

The condition for the existence of an unstable fixed point is that the real part of the eigenvalues of the Jacobian must be positive. If $\lambda$ is the eigenvalue, then the condition for oscillations is that $Re(\lambda) > 0$. The condition for oscillation for the dual regulation mechanism imposes a constrain on the slope of the regulatory functions, which is an augmentative combination of the conditions from the production and degradation regulation mechanisms:

$$-2\beta f(X^*) - \alpha g'(X^*) + \beta f'(X^*)X^* > 0 \tag{A.13}$$

This condition incorporates the conditions for oscillations from Eqn (A1.4) and from Eqn (A2.4). The ease of obtaining oscillations through the repression of production mechanism through scaling helps the condition for obtaining oscillations through the stimulation of degradation reaction, thereby making the dual regulation mechanism have the highest likelihood of obtaining oscillations.

**Appendix B: Generalized analysis for post-translational and transcriptional loop oscillator systems.**

In the main text of the paper, and in appendices A1-A3, we extensively discuss transcriptional loop oscillator systems, and the mechanisms of implementing the topology of such systems. However, the rules of design apply not just to transcriptional oscillators, but also to post-translational oscillator systems.

Transcriptional oscillators are systems where the production reaction has zeroth order kinetics, and degradation reaction has first order kinetics. In transcriptional oscillators, the number of molecules in the reactions is not fixed, and production-degradation events can change this number.

Post-translational oscillator systems (PTOs) on the other hand act as activator-deactivator systems, with a conserved number of molecules switching between active and inactive forms. Hence, the production and degradation kinetics in PTOs have first order kinetics. In this appendix, we demonstrate that the framework of Jacobian analysis that gives us the rules of design can be extended to include both types of systems.

##### 1. Condition for oscillations for generalized oscillator systems repression of production (RoP) or repression of activation (RoA)

Transcriptional oscillator dynamics can be written as:

$$\frac{dX}{dt} = \alpha g(Z) - \beta X \tag{B.1}$$

Similarly, post-translational oscillator dynamics, where the regulatory function g represses the activation reaction that switches $X$ from an inactive form to an active form can be written as:

$$\frac{dX}{dt} = \alpha(1-X)g(Z) - \beta X = \alpha g(Z) - (\beta + g(Z))X \tag{B.2}$$

The only difference between equations (B1.1) and (B1.2) is the extra term of $f(Z)$ in the degradation process in (B1.2). To combine the two types of systems, but still be able to distinguish the two types, we introduce a parameter into the equations.

$$\frac{dX}{dt} = \alpha g(Z) - (\beta + g(Z))X \tag{B.3}$$

where

$$\zeta = \begin{cases} 0 & \text{fortranscriptional oscillators} \\ 1 & \text{forpost-translational oscillators} \end{cases} \tag{B.4}$$

Now, we calculate the Jacobian as before, assuming a symmetric system with symmetric parameters and hence a symmetric fixed point at $fp = (X^*, X^*, X^*)$.

$$J_{(X^*,X^*,X^*)} = \begin{bmatrix} -\beta f(X^*) - \zeta\alpha g(X^*) & 0 & \alpha g'(X^*) - \beta f'(X^*)X^* - \zeta\alpha g'(X^*)X^* \\ \alpha g'(X^*) - \beta f'(X^*)X^* - \zeta\alpha g'(X^*)X^* & -\beta f(X^*) - \zeta\alpha g(X^*) & 0 \\ 0 & \alpha g'(X^*) - \beta f'(X^*)X^* - \zeta\alpha g'(X^*)X^* & -\beta f(X^*) - \zeta\alpha g(X^*) \end{bmatrix} \tag{B.5}$$

The condition for oscillations then becomes:

$$g'(X^*)(1-\zeta X^*) + 2\zeta g(X^*) < \frac{-2\beta}{\alpha} \tag{B.6}$$

In the case of $\zeta = 0$, the condition (B1.5) is the same as (A1.4). The post-translational RoA oscillator has a higher likelihood of sustaining oscillations due to the scaling of the regulatory function changing only the LHS of Eqn (B1.5), as in the case of the transcriptional oscillator.

##### 2. Condition for oscillations for generalized oscillator systems stimulation of degradation (SoD) or stimulation of deactivation (SoD)

Condition for oscillations for generalized transcriptional and post-translational oscillator systems with stimulation of degradation (SoD) or stimulation of deactivation (SoD) as the regulatory mechanism.

Transcriptional oscillator dynamics can be written as:

$$\frac{dX}{dt} = \alpha - \beta f(Z)X \tag{B.7}$$

Similarly, post-translational oscillator dynamics, where the regulatory function g represses the activation reaction that switches $X$ from an inactive form to an active form can be written as:

$$\frac{dX}{dt} = \alpha(1 - X) - \beta f(Z)X = \alpha - (\alpha + \beta f(Z))X \tag{B.8}$$

The only difference between equations (B2.1) and (B2.2) is the extra term of $\alpha$ in the degradation process in (B2.2). To combine the two types of systems, but still be able to distinguish the two types, we introduce a parameter into the equations.

$$\frac{dX}{dt} = \alpha - (\alpha\zeta + \beta f(Z))X \tag{B.9}$$

where,

$$\zeta = \begin{cases} 0 & \text{fortranscriptional oscillators} \\ 1 & \text{forpost-translational oscillators} \end{cases} \tag{B.10}$$

Now, we calculate the Jacobian as before, assuming a symmetric system with symmetric parameters and hence a symmetric fixed point at $fp = (X^*, X^*, X^*)$.

$$J_{(X^*,X^*,X^*)} = \begin{bmatrix} -\beta f(X^*) - \zeta\alpha & 0 & -\beta g'(X^*)X^* \\ -\beta g'(X^*)X^* & -\beta f(X^*) - \zeta\alpha & 0 \\ 0 & -\beta g'(X^*)X^* & -\beta f(X^*) - \zeta\alpha \end{bmatrix} \tag{B.11}$$

The condition for oscillations then becomes:

$$\beta g'(X^*) > \frac{2\beta g(X^*)}{X^*} + \frac{2\zeta}{\beta X^*} \tag{B.12}$$

In the case of $\zeta = 0$, the condition (B2.5) is the same as (A2.4). The post-translational SoD oscillator has a lower likelihood of sustaining oscillations as scaling changes both sides of Eqn (B2.5), as in the case o the transcriptional oscillator.

##### 3. Condition for oscillations for generalized oscillator systems with dual regulation

Condition for oscillations for generalized transcriptional and post-translational oscillator systems with dual regulation.

Transcriptional oscillator dynamics can be written as:

$$\frac{dX}{dt} = \alpha g(Z) - \beta f(Z)X \tag{B.13}$$

Similarly, post-translational oscillator dynamics, where the regulatory function g represses the activation reaction that switches X from an inactive form to an active form can be written as:

$$\frac{dX}{dt} = \alpha g(Z)(1 - X) - \beta f(Z)X = \alpha - (\alpha g(Z) + \beta f(Z))X \tag{B.14}$$

The only difference between equations (B3.1) and (B3.2) is the extra term of $\alpha f(Z)$ in the degradation process in (B3.2). To combine the two types of systems, but still be able to distinguish the two types, we introduce a parameter $\zeta$ into the equations.

$$\frac{dX}{dt} = \alpha - (\zeta\alpha g(Z) + \beta f(Z))X \tag{B.15}$$

where,

$$\zeta = \begin{cases} 0 & \text{fortranscriptional oscillators} \\ 1 & \text{forpost-translational oscillators} \end{cases} \tag{B.16}$$

Now, we calculate the Jacobian as before, assuming a symmetric system with symmetric parameters and hence a symmetric fixed point at $fp = (X^*, X^*, X^*)$.

$$J_{(X^*,X^*,X^*)} =$$
$$\begin{bmatrix} -\beta f(X^*) - \zeta\alpha g(X^*) & 0 & \alpha g'(X^*) - \beta f'(X^*)X^* - \zeta\alpha g'(X^*)X^* \\ \alpha g'(X^*) - \beta f'(X^*)X^* - \zeta\alpha g'(X^*)X^* & -\beta f(X^*) - \zeta\alpha g(X^*) & 0 \\ 0 & \alpha g'(X^*) - \beta f'(X^*)X^* - \zeta\alpha g'(X^*)X^* & -\beta f(X^*) - \zeta\alpha g(X^*) \end{bmatrix}$$

The condition for oscillations then becomes:

$$\frac{\beta f'(X^*)X^* - \alpha f'(X^*)(1 - \zeta X^*)}{2} > \beta f(X^*) + \zeta\alpha g(X^*) \tag{B.17}$$

In the case of $\zeta = 0$, the condition (B3.5) is the same as (A3.4). The post-translational dual oscillator has the highest likelihood of sustaining oscillations due to the co-operative combined effects of implementing repression through the regulation of both production and degradation mechanisms, as in the case of the transcriptional oscillator.

#### Appendix C: Nullcline analysis for incoherent input systems

We use nullcline analysis to analyze different possible regulatory implementations of the incoherent input system to predict the robustness response of a system. As an example, we will work through two cases to understand how to predict the optimal regulatory implementations for the incoherent input system.

**Principles of nullcline analysis:** The nullcline analysis rests on foundational biochemical notions. We use the fact that production rates can be as close to zero as possible, including being zero, which means that the regulation is allowed to be tight.

However, there is a non-zero lower bound on degradation rates $k_{min}^{deg}$. Even in the unrealistic case of zero rate of active degradation, there will be dilution as the cell grows, contributing to the lowering of the concentration of molecules in the cell.

Both production and degradation rates have upper bounds $k_{max}^{prod}$, and $k_{max}^{deg}$, from biophysical limits imposed due to the finite amount of resources available to cells, and the time it takes for the production and degradation machinery to assemble and carry out their function.

All parts of the biochemical phase space are not equally accessible. The biochemical zero is easily accessible, whereas a biochemical infinity is an unphysical limit. The notions of biochemically accessible phase space will implicate certain regulatory implementations as being easier - or harder - to achieve.

A nullcline profile that would support stable limit cycles would be as depicted in Figure 2A (or Equation Box 1 in main text).
The X nullcline has two extrema, and the region between the two extrema is called the switchback region. To obtain a stable limit cycle, the Y nullcline has to cross the X nullcline in the switchback region. To maximize the likelihood of such a crossing, we need to minimize $x_1$, maximize $x_2$, minimize $\Delta y$, and maximize $y_1$, as shown in SI.3. These conditions are not all independent of each other, and some conditions are implied by other conditions being satisfied. We consider different cases to analyze which regulatory mechanisms aid in achieving the necessary optimization of the nullclines.

##### 1. Determining the optimal regulatory implementations when Y is negatively regulated through the stimulation of the degradation rate

Determining the optimal regulatory implementations for the positive autoregulation of X and for the regulation of Y by X, when the negative regulation of X by Y is through the stimulation of the degradation reaction rate.

**Case I:** We consider each interaction, i.e. the YX, the XX, and the XY regulatory mechanisms to find the best possible implementation of each of these. As in the main text, S stands for Stimulation, R stands for Repression, P stands for Production, and D stands for Degradation. SoD, for instance, stands for Stimulation of Degradation.

**The YX interaction:** Assume that the YX interaction is fixed as SoD. We first identify the other X regulation that is most optimal.

**The XX interaction:** With the Y regulation on X fixed as SoD, we now need to identify the regulatory mechanism of X for positive auto-regulation between SoP and RoD. For the purposes of this calculation, we ignore dual interactions (the same analysis can be extended to dual mechanisms).

**X $\rightarrow$ X SoP:** If the XX interaction is assumed to be SoP, then the dynamical equation for X can be written as:

$$\frac{dX}{dt} = f(X) - \beta YX \tag{C.1}$$

Here, $f$ is a regulatory function that enhances the production rate, thereby representing the positive autoregulation of X. In the numerical results presented in the main text of the paper, we use Michaelis-Menten kinetics, and therefore a Hill function form for $f(X)$ as well as for the regulation of Y on X. However, for analytical tractability we use the linearized regulation for the Y interaction. The X nullcline can be obtained by setting the time derivative of X to zero:

$$Y = \frac{f(X)}{\beta X} \tag{C.2}$$

Although we use the Hill function form for our numerical results, analytically we can further generalize $f(X)$ to only include the asymptotic behavior. In the limit of small X values, we can write the asymptotic form of $f(X)$ to be:

$$\lim_{X \to 0} f(X) = a + bX^n \tag{C.3}$$

where a and b are constants related to the system parameters. We can see that the X nullcline blows up as $X \to 0$, but has a minimum at some value $X_{min}$. We find the extrema of the X nullcline as $X \to 0$.

$$\begin{aligned} &\frac{dY}{dX} = 0 \implies \frac{d}{dX}\left(\frac{f(X)}{\beta X}\right) = 0 \\ &\implies f'(X) = \frac{f(X)}{X} = bnX^{(n-1)} = \frac{a}{X} + bX^{(n-1)} \\ &\implies X_{min} = \left(\frac{a}{b(n-1)}\right)^{\frac{1}{n}} \end{aligned} \tag{C.4}$$

The functional form of the X nullcline in the large X region, the asymptote of X→ that enables a second turning point on the X nullcline is

$$\lim_{X \to \infty} f(X) = c - \frac{d}{X^m} \tag{C.5}$$

where c and d are constants related to the system parameters. We have the X-nullcline

$$Y = \frac{f(X)}{\beta X} \implies Y = c - \left[\frac{d}{X^m}\right]\frac{1}{\beta X} \tag{C.6}$$

Setting the Y derivative to zero, we obtain the local maximum point:

$$X_{max} = \left(\frac{d(m+1)}{c}\right)^{\frac{1}{m}} \tag{C.7}$$

To maximize the probability of the X and Y nullclines crossing, we need to minimize $x_1$, maximize $x_2$, minimize $\Delta y$, and maximize $y_1$.

**Minimize $x_1$:** This is equivalent to having $X_{min}$ as close to zero as possible. $X_{min}$ is given by:

$$X_{min} = \left(\frac{a}{b(n-1)}\right)^{\frac{1}{n}} \tag{C.8}$$

To minimize $x_1$, we need to have small values for a, and large values for b. Having small values of a is to have a tight regulation of the production rates, which is biochemically feasible.

**Maximize $y_1$:** To maximize $y_1$, we first calculate its value.

$$y_1 = Y(X_{min}) = \frac{f(X_{min})}{\beta X_{min}} = \frac{n}{\beta} a^{1-\frac{1}{n}} b^{\frac{1}{n}} (n-1)^{\frac{1}{n}-1} \tag{C.9}$$

Maximizing $y_1$ requires a and b to have large values, whereas we only have large value of b and small values of a from condition (C1.8). This constraint is not ideally supported by this regulatory mechanism, but only partially supported.

**Maximize $x_2$:** To maximize $x_2$, we consider the form of $X_{max}$.

$$X_{max} = \left[\frac{d(m+1)}{c}\right]^{\frac{1}{m}} \tag{C.10}$$

Maximizing d, and minimizing c will maximize $x_2$, and is compatible with the ease of accessibility of the biochemical phase space for these parameters.

**Minimize $\Delta y$:** To minimize $\Delta y$, we first calculate its functional form:

$$\begin{aligned}\Delta y &= Y(X_{max}) - Y(X_{min}) \\ \Delta y &= \left[mc^{\frac{(m+1)}{m}} d^{\frac{-1}{m}} (m+1)^{\frac{(-m-1)}{m}} - na^{\frac{(n-1)}{n}} b^{\frac{1}{n}} (n-1)^{\frac{(1-n)}{n}}\right] \frac{1}{\beta}\end{aligned} \tag{C.11}$$

To minimize Eqn (C1.11), having a small value for a is helpful, as well as having a small value for c. This constraint is compatible with the regulatory implementation considered here.

**X $\rightarrow$ X RoD:** If the XX interaction is assumed to be RoD, then the dynamical equation for X will be:

$$\frac{dX}{dt} = 1 - \beta Y g(X) X \tag{C.12}$$

Here, $g$ is a regulatory function that diminishes the degradation rate $\beta$ of X, thereby representing the auto-activation of X. In the numerical results presented in the main text of the paper, we use a Hill function form for $g(X)$ as well as for the regulation of Y on X. However, for analytical tractability we use the linearized regulation for the Y interaction. The X nullcline can be obtained by setting the time derivative of X to zero:

$$Y = \frac{1}{\beta g(X) X} \tag{C.13}$$

To have the X and Y nullclines cross, we need the same conditions as before, and the asymptotes of the repressive regulatory functions in different parts of the phase space have the same functional form. Specifically,

$$\lim_{X \to 0} \frac{1}{g(X)} \; a + bX^n \implies \lim_{X \to \infty} \frac{1}{g(X)} \; c - \frac{d}{X^m} \tag{C.14}$$

with small values for a and c, and large values for b and d. Therefore, the expressions for the maximum and minimum values of X, and y remain the same as in the case where the XX interaction was achieved through a stimulation of the production reaction.

**Minimize $x_1$:** This is equivalent to having $X_{min}$ as close to zero as possible. $X_{min}$ is given by:

$$X_{min} = \left[\frac{a}{b(n-1)}\right]^{\frac{1}{n}} \tag{C.15}$$

As before, it is advantageous to have small values of a, and large values of b. In this case, the inverse of the degradation rate corresponds to the physical interpretation of the parameter a. However, there is an upper bound on the degradation rate, i.e., $|\beta g(X)| < k^{deg}_{max}$ set by the degradation timescales in the system. This makes having small values of a unphysical, or at best biochemically hard to manage.

**Maximize $y_1$:** We know the expression for $y_1$ from Eqn (C1.9).

$$y_1 = Y(X_{min}) = \frac{f(X_{min})}{\beta X_{min}} = \left[\frac{n}{\beta a}\right]^{(1-\frac{1}{n})} b^{\frac{1}{n}}(n-1)^{(\frac{1}{n}-1)} \tag{C.16}$$

To maximize $y_1$ we need large values of a and b, which is better supported by this regulatory implementation of XX: RoD, compared to the implementation XX: SoP.

**Maximize $x_2$:** To maximize $x_2$, we consider the form of $X_{max}$.

$$X_{max} = \left[\frac{d(m+1)}{c}\right]^{\frac{1}{m}} \tag{C.17}$$

$x_2$ is maximized by having small values for c, and large values for the parameter d. However, as there is an upper bound on the degradation rates, and c is related to the inverse of the degradation rate constant of the system, $x_2$ can be maximized only within the bounds set by the degradation timescales. Thus, this condition is not well supported by the regulatory implementation of XX:RoD.

**Minimize $\Delta y$:** We know the expression for $\Delta y$ from Eqn (C1.11).

$$\begin{aligned}\Delta y &= Y(X_{max}) - Y(X_{min}) \\ \Delta y &= \left[mc^{\frac{(m+1)}{m}} d^{(\frac{-1}{m})}(m+1)^{(\frac{-m-1}{m})} - na^{(\frac{n-1}{n})}b^{(\frac{1}{n})}(n-1)^{(\frac{1-n}{n})}\right]\frac{1}{\beta}\end{aligned} \tag{C.18}$$

$\Delta y$ is minimized by having small values for a and c, which is not well accommodated by this regulatory implementation.

The conclusion from the comparison of the two XX implementations is that the XX: SoP regulatory implementation far better accommodates the biochemical and biophysical limits to support stable limit cycles than does the XX: RoD implementation.

**The XY interaction:** Now that we have fixed the regulation of X by Y as occurring through the stimulation of the degradation reaction, and the XX regulation as implemented through the stimulation of the production reaction, we need to deduce the final interaction in the system, which is the mechanism of regulation of Y by X.

The positive regulation of Y by X can be achieved either through the stimulation of the production reaction, or the through the repression of the degradation reaction.

**X → Y SoP:** If the XY interaction is assumed to be SoP, then the dynamical equation for Y can be written as:

$$\frac{dY}{dt} = f(X) - \beta Y \tag{C.19}$$

Here, $f$ is a regulatory function that enhances the production rate, thereby representing the positive regulation of Y by X. In the numerical results presented in the main text of the paper, we use Michaelis-Menten kinetics, and therefore a Hill function form for $f(X)$. The Y nullcline can be obtained by setting the time derivative of Y to zero to get the now familiar form of the nullcline:

$$Y = \frac{f(X)}{\beta} \tag{C.20}$$

Although we use the Hill function form for our numerical results, analytically we can further generalize $f(X)$ to only include the asymptotic behavior. In the limit of small X values, we can write the asymptotic form of $f(X)$ to be:

$$\lim_{X\to 0} f(X) = a + bX^n \tag{C.21}$$

where a and b are constants related to the system parameters. In the interest of maximizing the likelihood of the crossing of the X and the Y nullclines, we need to have the Y nullcline to start as close to the origin of the phase space as possible. This can be achieved by small values of a, which is the biochemical equivalent of having small production rates.

**X → Y RoD:** If the XY interaction is assumed to be RoD, then the dynamical equation for Y can be written as:

$$\frac{dY}{dt} = 1 - \beta Y g(X) \tag{C.22}$$

Here, $g$ is a regulatory function that diminishes the degradation rate $\beta$ of Y, thereby representing the positive regulation of Y by X. In the numerical results presented in the main text of the paper, we use a Hill function form for $g(X)$. The Y nullcline can be obtained by setting the time derivative of Y to zero:

$$Y = \frac{1}{\beta g(X)} \tag{C.23}$$

As before, we note that the Y nullcline is bounded below by the degradation rates of the system. As there is a maximum degradation rate $k^{deg}_{max}$, the Y nullcline cannot be as close to the x-axis as in the implementation of the XY interaction through SoP.

**Range of crossing:** The probability of obtaining stable limit cycles is directly proportional to the likelihood of the X and Y nullclines crossing in the switchback region. The switchback region is the box defined by $X_{min}$ and $X_{max}$ on the x-axis, and by $Y(X_{min})$ and $Y(X_{max})$ on the y-axis.

Here, we calculate the range of X values over which this crossing an occur. For a crossing to occur, the X nullcline and the Y nullclines must meet intersect at one point in phase space.

$$\begin{aligned} N_X = N_Y \implies \frac{a}{X} + \frac{bX^n}{X} &= \frac{c}{X} - \frac{d}{X^(m+1)} \\ \implies X_{cross} &= \left[\frac{(b+d)}{(c-a)}\right]^{\frac{1}{n}} \end{aligned} \tag{C.24}$$

We see that the range over which the crossing can occur increases as the separation between c and a decreases, as well as when the values of c and a are small. This is again, only possible to have in the XX: SoP, XY: SoP implementations when YX: SoD.

Hence, we conclude that the best possible implementation of the incoherent input system, given that the YX interaction is fixed to be YX: SoD is to have the XX interaction be XX: SoP, and the XY interaction be XY: SoP.

##### 2. Determining the optimal regulatory implementations when Y is negatively regulated through the repression of the production rate

Determining the optimal regulatory implementations for the positive autoregulation of X and for the regulation of Y by X, when the negative regulation of X by Y is through the repression of the production reaction rate.

**Case II:** We now consider the best implementation of the incoherent input system when the regulation of X by Y is through the repression of the production rate of X.

**The YX interaction:** Assume that the YX interaction is fixed as RoP. We first identify the other X regulation that is most optimal.

**The XX interaction:** With the Y regulation on X fixed as RoP, we now need to identify the regulatory mechanism of X for positive auto-regulation. The XX interaction can either be SoP or RoD.

**X $\rightarrow$ X SoP:** If the XX interaction is assumed to be SoP, then the dynamical equation for X can be written as:

$$\frac{dX}{dt} = f(X) + \alpha g(Y) - \beta X \tag{C.25}$$

$f$ is a regulatory function that enhances the production rates of X, thereby representing the positive autoregulation of X. Here, $g$ is a regulatory function that diminishes the production rate, thereby representing the negative regulation of X by Y. $\alpha$ is a constant that can be varied to encode the relative strengths of the regulations of the rate of production of X by X and Y.

In the numerical results presented in the main text of the paper, we use Michaelis-Menten kinetics, and therefore a Hill function form for $f(X)$ as well as for $g(Y)$. However, for analytical tractability we use the linearized forms of the regulatory functions. The linearized form of $g(Y)$ we use will be:

$$g(Y) \approx \frac{1}{(Y + K)} \tag{C.26}$$

The X nullcline can be obtained by setting the time derivative of X to zero:

$$Y = \alpha \left[\beta X - f(X)\right] - K \tag{C.27}$$

Although we use the Hill function form for our numerical results, analytically we can further generalize the regulatory functions to only include the asymptotic behavior. In the limit of small X values, we can write the asymptotic forms of $f(X)$ to be:

$$\lim_{X \to 0} f(X) = a + bX^n \tag{C.28}$$

Using (C2.4) in (C2.3), we get

$$Y(X) = \alpha \left[\beta X - a - bX^n\right] - K \tag{C.29}$$

In the limit of small X values, we have:

$$\lim_{X \to 0} Y(X) = -\alpha a - K \tag{C.30}$$

Since all the physical parameters of the system are positive, a negative value for the Y is disallowed. This implementation cannot support stable oscillations.

**X $\rightarrow$ X RoD:** If the XX interaction is assumed to be RoD, then the dynamical equation for X can be written as:

$$\frac{dX}{dt} = g(Y) - \beta g(X)X \tag{C.31}$$

Here, $g$ is a regulatory function that diminishes the production rate of X, thereby representing the negative regulation of Y on X. The function $g$ also diminishes the degradation rate of X, thereby representing the positive autoregulation of X.

In the numerical results presented in the main text of the paper, we use Michaelis-Menten kinetics, and therefore a Hill function form for $g$. However, for analytical tractability we use the linearized regulation for the Y interaction. The linearized form of $g(Y)$ we use will be:

$$g(Y) \approx \frac{1}{Y + K} \tag{C.32}$$

The X nullcline can be obtained by setting the X derivative to zero:

$$Y = \frac{1}{\beta g(X)X} - K \tag{C.33}$$

As in previous cases when we encountered nullclines of this form, we conclude that the likelihood of obtaining oscillations are low as the X nullcline is constrained by the upper bound set by the degradation rates of the system.

**The XY interaction:** The regulation of Y be X was analyzed independently of the regulation of X by Y and the positive autoregulation of X. The results from Eqns (C1.19)-(C1.23) still hold.

In conclusion, we note that oscillations are either not supported at all, or supported only weakly when the negative regulation of X through Y is implemented through the repression of the production rates.

#### Appendix D: Nullcline analysis for coherent input systems

Using the same biochemical principles from Appendix C, we now deduce the optimal regulatory implementations for the coherent input systems for one case, when the positive regulation of X by Y is through the repression of the degradation reaction.

**Case I:** We consider each interaction, i.e. the YX, the XX, and the XY regulatory mechanisms to find the best possible implementation of each of these. As in the main text, S stands for Stimulation, R stands for Repression, P stands for Production, and D stands for Degradation. So, SoD stands for Stimulation of Degradation, for example.

**The YX interaction:** Assume that the YX interaction is fixed as RoD. We first identify the other X regulation that is most optimal.

**The XX interaction:** With the Y regulation on X fixed as RoD, we now need to identify the regulatory mechanism of X for positive auto-regulation. For the purposes of this calculation, we ignore dual interactions (the same analysis can be extended to include dual mechanisms). The XX interaction can either be SoP or RoD.

**X → X SoP:** If the XX interaction is assumed to be SoP, then the dynamical equation for X can be written as:

$$\frac{dX}{dt} = f(X) - \beta g(Y)X \tag{D.1}$$

Here, $f$ is a regulatory function that enhances the production rate, thereby representing the positive autoregulation of X. $g$ is a regulatory function that diminishes the degradation rate $\beta$ of X. In the numerical results presented in the main text of the paper, we use Michaelis-Menten kinetics, and therefore a Hill function form for $f(X)$ as well as for $g(Y)$. However, for analytical tractability we use the linearized regulation for the Y interaction. The linearized form of $g(Y)$ we use will be:

$$g(Y) \approx \frac{1}{Y+K} \tag{D.2}$$

The X nullcline can be obtained by setting the time derivative of X to zero:

$$Y = \frac{\beta X}{f(X)} - K \tag{D.3}$$

Although we use the Hill function form for our numerical results, analytically we can further generalize $f(X)$ to only include the asymptotic behavior. In the limit of small X values, we can write the asymptotic form of $f(X)$ to be:

$$\lim_{X}$$

$\rightarrow$ 0 f(X) = a + b X$^n$(D.4)
where a and b are constants related to the system parameters. We can see that the X nullcline blows up as $X \rightarrow 0$, but has a minimum at some value $X_{min}$. We find the extrema of the X nullcline as $X \rightarrow 0$.

$$\begin{aligned} \frac{dY}{dX} = 0 &\implies \frac{d}{dX}\left(\frac{\beta X}{f(X)} - K\right) = 0 \\ &\implies f'(X) = \frac{f(X)}{X} = bnX^{(n-1)} = \frac{a}{X} + bX^{(n-1)} \\ &\implies X_{min} = \left[\frac{a}{b(n-1)}\right]^{\left(\frac{1}{n}\right)} \end{aligned} \tag{D.5}$$

where a and b are constants related to the system parameters.

The functional form of the X nullcline in the large X region, the asymptote of $X \rightarrow \infty$ that enables a second turning point on the X nullcline is:

$$\lim_{x \rightarrow \infty} f(X) = c - \frac{d}{X^m} \tag{D.6}$$

where c and d are constants related to the system parameters. We have the X-nullcline

$$Y = \frac{\beta X}{f(X)} - K \implies Y = \frac{\beta X}{\left[c - \frac{d}{X^m}\right]} - K \tag{D.7}$$

Setting the Y derivative to zero, we obtain the local maximum point:

$$X_{max} = \left[\frac{d(m+1)}{c}\right]^{\left(\frac{1}{m}\right)} \tag{D.8}$$

To maximize the probability of the X and Y nullclines crossing, we need to minimize $x_1$, maximize $x_2$, minimize $\Delta y$, and maximize $y_1$.

**Minimize $x_1$:** This is equivalent to having X$_m inasclosetozeroaspossible.$X$_{min}$ is given by:

$$X_{min} = \left[\frac{a}{b(n-1)}\right]^{\left(\frac{1}{n}\right)} \tag{D.9}$$

To minimize $x_1$, we need to have small values for a, and large values for b. Having small values of a is to have a tight regulation of the production rates, which is biochemically feasible.

**Maximize $y_1$:** To maximize $y_1$, we first calculate its value.

$$y_1 = Y(X_{min}) = \frac{\beta X_{min}}{f(X_{min})} - K = \left(\frac{\beta}{n}a\right)^{(\frac{1}{n}-1)} (b)^{(\frac{-1}{n})} (n-1)^{(1-\frac{1}{n})} - K \tag{D.10}$$

Maximizing $y_1$ requires a and b to have large values, whereas we only have large value of b and small values of a from condition (C1.8). This constraint is not ideally supported by this regulatory mechanism, but only partially supported.

**Maximize $x_2$:** To maximize $x_2$, we consider the form of $X_{max}$.

$$X_{max} = \left[\frac{d(m+1)}{c}\right]^{(\frac{1}{m})} \tag{D.11}$$

Maximizing d, and minimizing c will maximize $x_2$, and is compatible with the ease of accessibility of the biochemical phase space for these parameters.

**Minimize $\Delta$ y:** To minimize $\Delta y$, we first calculate its functional form:

$$\begin{aligned} \Delta y &= Y(X_{max}) - Y(X_{min}) \\ \Delta y &= \beta \left[\frac{1}{m} c^{\frac{(-m+1)}{m}} d^{\frac{1}{m}} (m+1)^{\frac{(m+1)}{m}}\right)^{-\frac{1}{n}} a^{\frac{(1-n)}{n}}\Big) b^{(\frac{-1}{n})} (n-1)^{\frac{(n-1)}{n}} \right] \end{aligned} \tag{D.12}$$

To minimize Eqn (C1.11), having a small value for a is helpful, as well as having a small value for c. This constraint is compatible with the regulatory implementation considered here.

**X $\rightarrow$ X RoD:** If the XX interaction is assumed to be RoD, then the dynamical equation for X can be written as:

$$\frac{dX}{dt} = 1 - g(X)X - \beta g(Y)X \tag{D.13}$$

Here, $g$ is a regulatory function that diminishes the degradation rate $\beta$ of X, thereby representing the positive regulation of X by Y. The autoactivation of X is represented by the regulation of the degradation rate of X. The parameter $\beta$ denotes the relative rates of degradation between the autoactivation and the positive regulation of X by Y.

In the numerical results presented in the main text of the paper, we use Michaelis-Menten kinetics, and therefore a Hill function form for $g(X)$ as well as for the regulation of Y on X. However, for analytical tractability we use the linearized regulation for the Y interaction. The linearized form of $g(Y)$ we use will be:

$$g(Y) \approx \frac{1}{(Y+K)} \tag{D.14}$$

The X nullcline can be obtained by setting the time derivative of X to zero:

$$Y = \frac{betaX}{1 + \alpha g(X)X} - K \tag{D.15}$$

At small X values, the Y nullcline tends to negative values, which is unphysical, making this implementation less likely to support stable limit cycles.

**The XY interaction:** Now that we have fixed the regulation of X by Y as occurring through the repression of degradation reaction, and the XX regulation implemented through the stimulation of the production reaction, we need to deduce the final interaction in the system, which is the mechanism of regulation of Y by X.

The negative regulation of Y by X can be achieved either through the repression of the production reaction, or the through the stimulation of the degradation reaction, or both. Here we consider all three cases to show that the dual regulation helps the robustness response of coherent input systems.

**X → Y RoP:** If the XY interaction is assumed to be RoP, then the dynamical equation for Y can be written as:

$$\frac{dY}{dt} = g(X) - \beta Y \tag{D.16}$$

Here, $g$ is a regulatory function that diminishes the production rate, thereby representing the negative regulation of Y by X. In the numerical results presented in the main text of the paper, we use Michaelis-Menten kinetics, and therefore a Hill function form for $g(X)$. The Y nullcline can be obtained by setting the time derivative of Y to zero to get the nullcline:

$$Y = \frac{g(X)}{\beta} \tag{D.17}$$

$g$ is a saturating, monophasic function. There is an upper bound on the degradation rate $\beta$, and there is a lower bound on the degradation rate, set by the dilution factor due to cell growth. Hence, the Y nullcline is bound between two values, shrinking the range of values where crossings with the X nullcline can occur.

**X →Y SoD:** If the XY interaction is assumed to be SoD, then the dynamical equation for Y can be written as:

$$\frac{dY}{dt} = 1 - \beta f(X) Y \tag{D.18}$$

Here, $f$ is a regulatory function that enhances the degradation rate $\beta$ of Y. The Y nullcline can be obtained by setting the time derivative of Y to zero:

$$Y = \frac{1}{\beta f(X)} \tag{D.19}$$

This function falls off quickly, and can cross the X nullcline in the switchback region.

**X →Y RoP  SoD:** If the XY interaction is assumed to be RoP  SoD, then the dynamical equation for Y can be written as:

$$\frac{dY}{dt} = g(X) - \beta f(X) Y \tag{D.20}$$

Here, $f$ is a regulatory function that enhances the degradation rate $\beta$ of Y, and $g$ is a regulatory function that diminishes the production rate of Y. The Y nullcline can be obtained by setting the time derivative of Y to zero:

$$Y = \frac{g(X)}{\beta f(X)} (D.21) \tag{D.21}$$

Having regulations on both production and degradation steps helps the system. The function $g(X)$ has small values when X is small, and the function inverse $f(X)$ has extremely high values when X is small, but falls off quickly. With the two regulations in place, the function $f(X)$ is modulated by the low values of $g(X)$ to cross the X nullcline closer to the minimum value of the X nullcline, making the likelihood of crossing within the switchback region higher. We see that the dual regulation for the XY interaction is a preferred regulatory implementation in the three most robust implementations of the coherent input system, for this reason.

**A note on the differential functions of dual regulation in coherent and incoherent input systems:**

Seeing how the dual regulatory implementations aid in increasing the likelihood of the X and Y nullclines crossing, we ask why it is that we do not observe dual implementations leading to robust responses in the most robust regulatory implementation of the incoherent input system.

To understand why the dual implementation might not be as beneficial in the incoherent input system, let us consider dual regulation of X by Y, i.e. YX: RoP  SoD, XX: SoP. The dynamical equation for X gives us:

$$\frac{dX}{dt} = f(X) + \alpha g(Y) - \beta YX \tag{D.22}$$

Here, $f$ is a regulatory function that enhances the production rate of X, indicating the positive autoregulation. $g$ is a regulatory function that diminishes the production rate of X, through Y, representing the negative regulation of X by Y, via repression of production rate $\alpha$, and the negative regulation by Y on X through the stimulation of the degradation rate $\beta$ is expressed as a linear action. Setting the time derivative of X to zero, we get the X nullcline:

$$f(X) + \alpha g(Y) = \beta YX \tag{D.23}$$

Using the linearized form of $g$, we have:

$$Y = \frac{\alpha g(Y)}{\beta X} + \frac{f(X)}{\beta X} \tag{D.24}$$

The additional term from the dual regulation doesn't change the nullcline properties by a lot, and hence isn't beneficial to incoherent input systems.

#### Appendix E: Nullcline analysis of the mammalian circadian oscillator

The mammalian oscillator consists of positive and negative feedback arms which function together to give stable 24 hour oscillations. BMAL1 and CLOCK proteins make a hetero-dimer, which then binds to the E-box promoters of PER and CRY genes, and activate their transcription through SoP.

Per and Cry proteins, in turn form a hetero-dimer complex, inhibiting the activity of BMAL1-CLK complex through SoD. Finally, an increased concentration of PER in the nucleus mediates the up-regulation of BMAL1 transcription, and thus BMAL1 complex achieves positive autoregulation through the stimulation of its own production by SoP.

The other important component of the circadian oscillations is the negative autoregulation by BMAL1. We determine the regulatory implementation of this negative interaction through the principles laid out above. The negative interaction can be achieved through RoP, or SoD. Here, we outline the constraints to get the largest ranges of X values with the two types of implementation. X denotes the BMAL1-CLK complex, Y denotes the PER-CRY complex.

**Case I:** Negative autoregulation achieved through RoP

$$\frac{dX}{dt} = \alpha_1 f(X) + \alpha_2 g(X) - \beta YX \tag{E.1}$$

For the X nullcline, we set the derivative of X to zero, which gives:

$$\frac{\alpha_1 f(X) + \alpha_2 g(X)}{\beta X} = Y \tag{E.2}$$

To get the extremum value of X, we set $dY/dX$ to zero:

$$\frac{dY}{dX} = 0 \implies X^n = \frac{\alpha_1 K^n}{nK^n(\alpha_2 - \alpha_1) - \alpha_2} \tag{E.3}$$

To avoid unphysical values of concentrations of X, we have the constraint that

$$nK^n(\alpha_2 - \alpha_1) - \alpha_2 > 0 \implies \frac{\alpha_1}{\alpha_2} < 1 - \frac{1}{nK^n} \tag{E.4}$$

This constraint implies that the as long as the rate of production of X is not repressed to the extent that the concentration of X is too low, the system can support oscillations.

**Case II:** Negative autoregulation is achieved through SoD

$$\frac{dX}{dt} = \alpha g(X) - \beta_1 f(X)X - \beta_2 YX \tag{E.5}$$

For the X nullcline, we set the derivative of X to zero, which gives:

$$Y = \frac{\alpha g(X) - \beta_1 f(X)X}{\beta_2 X} \tag{E.6}$$

To get the extremum value of X, we set $\frac{dY}{dX}$ to zero:

$$\frac{dY}{dX} = \frac{\alpha g'(X)X - \alpha g(X)}{\beta_2 X^2} = \frac{\beta_1}{\beta_2} f'(X) \implies X^{(n+1)} + \frac{\alpha}{\beta n}(n-1)X^n - \frac{\alpha K^n}{\beta n} s = 0 \tag{E.7}$$

This constraint has to be satisfied to get oscillations in the system. This is a harder criterion to satisfy, compared to Case I.

#### APPENDIX SUPPLEMENTARY METHODS

##### M1. Convergence Criteria for limit cycles

To determine whether the oscillations converged to a stable limit cycle, we use the fact that a deterministic system has near-perfect autocorrelation for undamped oscillations. We run the ODEs time t such that $t \gg T$, where T is the period of oscillation for that parameter set. t is set to be sufficient for at least 8 cycles of oscillations, and if the height of the last peak of the oscillation is within 1% of the first peak, then we say that a stable limit cycle is reached.

##### M2. Calculation of Phase Coherence Values

Phase-coherence of oscillations are calculated in the stochastic limit when the system size is finite. We use the Gillespie Stochastic Simulation Algorithm (SSA) to calculate the time series for each parameter set. To ensure convergence of phase coherence values for each time series, or equivalently, to establish an error bound on the calculated values of phase coherence of oscillations, we follow these steps for each parameter set:

i) Run the ODEs for some time t such that $t \gg T$, where T is the period of oscillation for that parameter set.

ii) For each dynamical variable, calculate the autocorrelation function of the time series, and note the position $(p_1, p_2)$ and heights $(h_1, h_2)$ of the first two non-unity peak of the autocorrelation function.

iii) Run the system again for a time t, and calculate the positions $(p_3, p_4)$ and heights $(h_3, h_4)$ of the first two non-unity peaks of the autocorrelation function.

iv) The system is said to reach convergence when the first non-unit peak position and heights from the two series ($p_1, p_3$ and $h_1, h_3$) are aligned to within 0.001% of each other for each dynamical variable in the system.

v) The height of the first non-unity peak of the longer time series $(h_3)$ is the phase-coherence value of the oscillation for the parameter set, for that dynamical variable and iteration.

vi) We repeat the steps (i)-(v) over 50 iterations of the Gillespie SSA, and calculate the phase coherence values for each iteration, and note the phase coherence values for each variable.

vii) Finally, we average over the phase coherence values from the 50 iterations of the Gillespie algorithm, and over the phase coherence values of all dynamical variables in the system to get the calculated average phase coherence values that are used for further analyses.

#### M3. Positive-Negative Oscillator Motifs

We describe the parameter ranges used for the calculation of robustness, and for the calculation of phase coherence values in the positive-negative oscillator motifs. The equations used to model different systems are based on the interactions at the ode, and the regulatory implementations. We demonstrate the general functional forms used here.

##### M3.1. Gene Expression Networks with Positive-Negative Oscillator Motifs

Each system has basal production $(k_{aX}, k_{aY})$ and degradation $(k_{dX}X, k_{dYY})$ terms.

- Stimulation of Y by X through SoP is modelled as $k_{XY}\frac{X^n}{(K^n+X^n)}$
- Stimulation of Y by X through RoD is modelled as $-k_{XY}\frac{K^n}{K^n+X^n}Y$
- Repression of X by Y through SoD is modelled as $-k_{YX}\frac{K^n}{K^n+Y^n}X$
- Repression of X by Y through RoP is modelled as $k_{YX}\frac{K^n}{K^n+Y^n}$
- Positive autoregulation of X through SoP is modelled as $k_{XX}\frac{X^n}{K^n+X^n}$
- Positive autoregulation of X through RoD is modelled as $-k_{XX}\frac{K^n}{K^n+X^n}X$

The distributions of the parameters are:

| System Parameter | Ranges Used |
|---|---|
| $\alpha_{0X}, \alpha_{0Y}$ | $[0.001, 0.01]\mu M min^{-1}$ |
| $\beta_{0X}, \beta_{0Y}$ | $[0.001, 0.01]$ min$^{-1}$ |
| $k$ | $[10^{-3}, 10^{2}]$ min$^{-1}$ |
| $K$ | $[10^{-3}, 10^{2}]$ $\mu$M |
| $n$ | $[1, 4]$ |

**Appendix Table S1:** Parameter ranges used for transcriptional positive-negative oscillator systems. Here, k represents all other rate constants apart from the basal production and degradation rates.

The 27 implementations for the incoherent, and coherent input systems produce distinct responses in terms of the fraction of parameter sets which support stable limit cycles, as shown in Figure 4.

##### M3.2. Post-translational Oscillators with Positive-Negative Oscillator Motifs

We describe the parameter ranges used for the calculation of robustness, and for the calculation of phase coherence values in the positive-negative oscillator motifs. The equations used to model different systems are based on the interactions at the ode, and the regulatory implementations. We demonstrate the general functional forms used here.

- Each system has basal production ($k_{aX}(1-X), k_{aY}(1-Y)$) and degradation ($k_{dX}X, k_{dY}Y$) terms.
- Stimulation of Y by X through SoP is modelled as $k_{XY}\frac{X^n}{K^n+X^n}(1-Y)$
- Stimulation of Y by X through RoD is modelled as $-k_{XY}\frac{K^n}{K^n+X^n}Y$
- Repression of X by Y through SoD is modelled as $-k_{YX}\frac{K^n}{K^n+Y^n}X$
- Repression of X by Y through RoP is modelled as $k_{YX}\frac{K^n}{K^n+Y^n}(1-X)$
- Positive autoregulation of X through SoP is modelled as $k_{XX}\frac{X^n}{K^n+X^n}(1-X)$
- Positive autoregulation of X through RoD is modelled as $-k_{XX}\frac{K^n}{K^n+X^n}X$

The distributions of the parameters are:

| System Parameter | Ranges Used |
|---|---|
| $\alpha_{0X}, \alpha_{0Y}$ | $[0.001, 0.01]\mu Mmin^{-1}$ |
| $\beta_{0X}, \beta_{0Y}$ | $[0.001, 0.01]$ min$^{-1}$ |
| $k$ | $[10^{-3}, 10^{2}]$ min$^{-1}$ |
| $K$ | $[10^{-3}, 10^{2}]$ $\mu$ M |
| $n$ | $[1, 4]$ |

**Appendix Table S2:** Parameter ranges used for post-translational positive-negative oscillator systems. Here, k represents all other rate constants apart from the basal production and degradation rates.

We see that there are preferred regulatory implementations in positive-negative PTO systems, just as in the case of transcriptional positive-negative oscillator systems.

##### M4. Parameter range determination for Loop Oscillators

As described in the main text, the negative feedback loop of the repressilator can be modelled in three different ways: through the repression of production the stimulation of degradation, and through dual regulation. For each implementation of the repressilator loop, we explore a large range of parameters within the experimentally established limits of parameters observed in a bacterial cell.

For each of the three implementations, we then consider only those regions of parameter space that support stable limit cycle dynamics for further numerical analyses.

###### M4.1. Parameter range determination for Gene Expression Networks (GENs) of Loop Oscillators

The mathematical representation of each implementation is shown in Appendix Table S3. Here, $\alpha$'s correspond to the unregulated production rates, $\beta$'s correspond to the unregulated degradation rates, K's correspond to the affinity parameters of the system, and n's correspond to the Hill coefficients.

| Repression of Production | Stimulation of Degradation | Dual Regulation |
|---|---|---|
| $\frac{dX}{dt} = \alpha_1 \frac{K_1^{n1}}{K_1^{n1}+Z^{n1}} - \beta_1 X$ | $\frac{dX}{dt} = \alpha_1 - \beta_1 \frac{Z^{n1}}{K_1^{n1}+Z^{n1}} X$ | $\frac{dX}{dt} = \alpha_1 \frac{K_a 1^{(}n_1))}{K_{a1}^{n1}+Z^{n1}} - \beta_1 \frac{Z^{n2}}{K_{d1}^{n2}+Z^{n2}} X$ |
| $\frac{dY}{dt} = \alpha_2 \frac{K_2^{n2}}{K_2^{n2}+X^{n2}} - \beta_2 Y$ | $\frac{dY}{dt} = \alpha_2 - \beta_2 fracX^{n2} K_2^{n2} + X^{n2} Y$ | $\frac{dY}{dt} = \alpha_2 \frac{K_{a2}^{n3}}{K_{a2}^{n3}+X^{n3}} - \beta_2 \frac{X^{(}n_4)}{K_{d2}^{n4}+X^{n4}} Y$ |
| $\frac{dZ}{dt} = \alpha_3 \frac{(K_3^{n3}}{K_3^{n3}+Y^{n3}} - \beta_3 Z$ | $\frac{dZ}{dt} = \alpha_3 - \beta_3 \frac{Y^{n3}}{K_3^{n3}+Y^{n3}} Z$ | $\frac{dZ}{dt} = \alpha_3 \frac{(K_{a3}^{n5}}{K_{a3}^{n5}+Y^{n5}} - \beta_3 \frac{Y^{n6}}{K_{d3}^{n6}+Y^{n6}} Z$ |

**Appendix Table S3:** System of differential equations for the asymmetric repressilator system, where the production is a zeroth kinetic order process, and the degradation is a first kinetic order process. Column I shows the negative loop modelled as the repression of the production rates. Column II shows the negative loop modelled as the stimulation of the degradation rates. Column III shows the negative loop modelled as Dual regulation.

The typical rates for the production of proteins is experimentally determined as being in the range of a few molecules per min [21]. The distribution we use for the exploration of rates are as follows:

| System Parameter | Ranges Used |
|---|---|
| $\alpha$ | $[10^{-2}, 10^{2}] \mu M min^{-1}$ |
| $\beta$ | $[10^{-3}, 0.2]$ min$^{-1}$ |
| $K_a, K_d, K$ | $[1, 10^{3}]$ $\mu M$ |
| $n$ | $[4, 5]$ |

**Appendix Table S4:**Parameter range determination for transcriptional loop oscillators.

To determine the ranges of parameter distributions to use for numerical calculations, we first swept parameters over orders of magnitude to narrow down regions of interest, where the parameter values fall within the range of experimentally observed parameter values for the systems, and which support stable limit cycle oscillations, as shown in Fig. S3.

##### M4.2. Parameter range determination for Post-Translational Loop Oscillators (PTOs)

The mathematical representation of each implementation is shown in Appendix Table S5. Here, $\alpha$'s correspond to the unregulated production rates, $\beta$'s correspond to the unregulated degradation rates, $K$'s correspond to the affinity parameters of the system, and n's correspond to the Hill coefficients.

| Repression of Production | Stimulation of Degradation | Dual Regulation |
|---|---|---|
| $\frac{dX}{dt} = \alpha_1(1-X)\frac{(K_1^{n_1}}{K_1^{n_1}+Z^{n_1}} - \beta_1 X$ | $\frac{dX}{dt} = \alpha_1(1-X) - \beta_1 \frac{Z^{n_1}}{K_1^{n_1}+Z^{n_1}} X$ | $\frac{dX}{dt} = \alpha_1(1-X)\frac{(K_{a1}^{n_1}}{K_{a1}^{n_1}+Z^{n_1}} -_1 Z^(n_2)/(K_d 1^(n_2) + Z^(n_2))X$ |
| $\frac{dY}{dt} = \alpha_2(1-Y)\frac{K_2^{n_2}}{K_2^{n_2}+X^{n_2}} - \beta_2 Y$ | $\frac{dY}{dt} = \alpha_2(1-Y) - \beta_2 \frac{X^{n_2}}{K_2^{n_2}+X^{n_2}} Y$ | $\frac{dY}{dt} = \alpha_2(1-Y)\frac{K_{a2}^{n_3}}{K_{a2}^{n_3}+X^{n_3}} - \beta_2 \frac{X^{n_4}}{K_{d2}^{n_4}+X^{n_4}} Y$ |
| $\frac{dZ}{dt} = \alpha_3(1-Z)\frac{(K_3^(n_3))}{(K_3^(n_3)+Y^(n_3))} -_3 Z$ | $\frac{dZ}{dt} = \alpha_3(1-Z) - \beta_3 \frac{Y^{n_3}}{K_3^{n_3}+Y^(n_3} Z$ | $\frac{dZ}{dt} = \alpha_3(1-Z)\frac{K_{a3}^{n_5}}{K_{a3}^{n_5}+Y^{n_5}} - \beta_3 \frac{Y^{n_6}}{K_{d3}^{n_6}+Y^{n_6}} Z$ |

**Appendix Table S5:** System of differential equations for the asymmetric repressilator system, where the production is a zeroth kinetic order process, and the degradation is a first kinetic order process. Column I shows the negative loop modelled as the repression of the production rates. Column II shows the negative loop modelled as the stimulation of the degradation rates. Column III shows the negative loop modelled as Dual regulation.

We see that in the PTOs, where the number of molecules is conserved, i.e. there is no production or degradation of molecules, but the molecules switch from active to inactive forms, the same trend of obtaining oscillations with higher probability for repressilator loop executed through the repression of the activation compared to the repression through the stimulation of the deactivation. As in GENs, we see that dual regulation is the most robust implementation of the negative feedback.

The parameter sets used for the three implementations are shown in the table below.

| System Parameter | Range for RoA | Range for SoD | Range for Dual Regulation |
|---|---|---|---|
| $\alpha$ | $1\mu$ M min$^{-1}$ | $[10^{-6}, 0.12]\ \mu M min^{-1}$ | $1\mu$ M min$^{-1}$ |
| $\beta$ | $[10^{-4}, 0.5]$ min$^{-1}$ | 1 min$^{-1}$ | 1 min$^{-1}$ |
| $K$ | $[10^{-4}, 0.5]\ \mu M$ | $[10^{-6}, 2]\ \mu M$ | - |
| $K_a$ | - | - | $[10^{-8}, 0.5]\ \mu M$ |
| $K_d$ | - | - | $[10^{-8}, 2]\ \mu M$ |
| $n$ | $[4, 5]$ | $[4, 5]$ | $[4, 5]$ |

**Appendix Table S6:**Parameter ranges used for post-translational loop oscillators .

The negative loop of the repressilator motif, when implemented via dual regulation is the most robust implementation in PTO systems, where the product and degradation are first kinetic order processes (see Fig S5).

Implementation of the negative regulation through repressing the activation rate is more robust than implementing the negative regulation by stimulating the deactivation rate. We see that the post translational repressilator motif follows the trends seen in transcriptional repressilator system.

#### M5. Phase Coherence of Symmetric and Asymmetric Post-translational Repressilator Motifs

- To calculate the robustness response of each implementation of the repressilator, we draw parameter sets at random from the uniform random distributions of parameters described in Table M6 until we have 500 parameter sets that give stable limit cycles.

- To determine that we have converged into a stable state, we use convergence criteria mentioned in Methods M1. Robustness is then defined as 500 over the number of parameter sets that were drawn from the distributions to get 500 sets that give stable oscillations.

- For the parameter sets that support stable limit cycles, we calculate the phase coherence values using the procedure in E.

- We repeat this process for symmetric and asymmetric PTO repressilator motifs.

Fig S5 shows the results of the steps M5 (i)-(iv). As in the case of the transcriptional repressilator system, we see that the symmetric systems saturate the bound on the maximum phase coherence values that can be achieved at a given deterministic arclength, and a given system size $\Omega$. Here, the system size also corresponds to the number of molecules of active forms of each interacting gene product in the system.

#### M6. Phase Coherence of Symmetric and Asymmetric Post-translational Repressilator Motifs

| System Parameter | Ranges Used |
|---|---|
| $\alpha_0, \beta_0$ | $[0.001, 0.01]$ $\mu$ M min$^{-1}$, min$^{-1}$ |
| $\alpha, \beta$ | $\left[10^{-3}, 10^{2}\right]$ $\mu$ min$^{-1}$, min$^{-1}$ |

**Appendix Table S7:** Experimentally motivated distributions of parameter from which the function parameters are drawn.

SoP:

$$\frac{dY}{dt} = \alpha_0 + \alpha f(X) - \beta_0 Y = 0$$

$$\implies Y = \frac{\alpha_0}{\beta_0} + \frac{\alpha}{\beta_0} f(X)$$

$$Y \approx \frac{\alpha_0}{\beta_0} + \frac{\alpha}{\beta_0}\left[f(X_{mid}) + f'(X_{mid})(X - X_{mid})\right] \tag{E.8}$$

Assuming $X_{mid} \approx K$,

$$Y \approx \frac{\alpha_0}{\beta_0} + \frac{\alpha}{\beta_0}\left[\frac{1}{2} + \frac{n}{4K(X-K)}\right]$$

$$\text{Slope} = \frac{}{0}\frac{n}{4K}$$

$$\text{Range of possible slopes: } \left[10^{-1}, 10^{5}\right]\frac{n}{4K} \tag{E.9}$$

RoD:

$$\frac{dY}{dt} = \alpha_0 - \beta_0 Y - \beta g(X) Y = 0$$

$$\implies Y = \frac{\alpha_0}{\beta_0 + \beta g(X)}$$

$$Y \approx \frac{\alpha_0}{\beta_0 + \beta\left[g(X_{mid}) + g'(X_{mid})(X - X_{mid})\right]} \tag{E.10}$$

Assuming $X_{mid} \approx K$,

$$Y \approx \frac{\alpha_0}{(\beta_0 + \beta[\frac{1}{2} - \frac{n}{4K}(X-K)])} \approx \frac{\alpha_0}{(C - bX)}$$

$$Y \ \alpha_0[1 + \frac{b}{C}X]$$

$$\text{where} C = \beta_0 + \frac{}{2}[1 + \frac{n}{2}]; b = \alpha_0\beta \tag{E.11}$$

If $n = 4$,

$$\text{Slope} = \frac{(\alpha_0\beta)}{(\beta_0 + 1.5\beta)}\frac{n}{4K}$$

$$\text{Range of possible slopes: } \left[10^{-4}, 400\right]\frac{n}{4K} \tag{E.12}$$

#### APPENDIX SUPPLEMENTARY FIGURES

##### Fig S1. Performance of 27 implementations of post-transcriptional positive-negative oscillators

The ranked response for the fraction of parameter sets supporting stable limit cycles in post-translational oscillator systems is shown below.

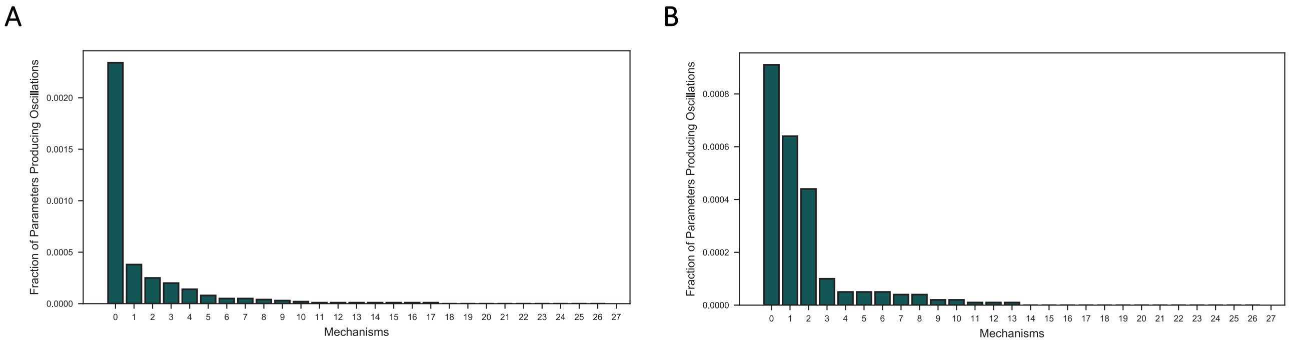


**Fig S1:** The bar graphs show the fraction of parameter sets that support stable limit cycles for the post-transcriptional positive-negative oscillator systems. The panel on the left shows the results for the 27 implementations of the incoherent input system, and the panel on the right shows the results for the 27 implementations of the coherent input system.

**Fig S2. Distributions of the nullcline parameters in two implementations of the X and Y regulations**

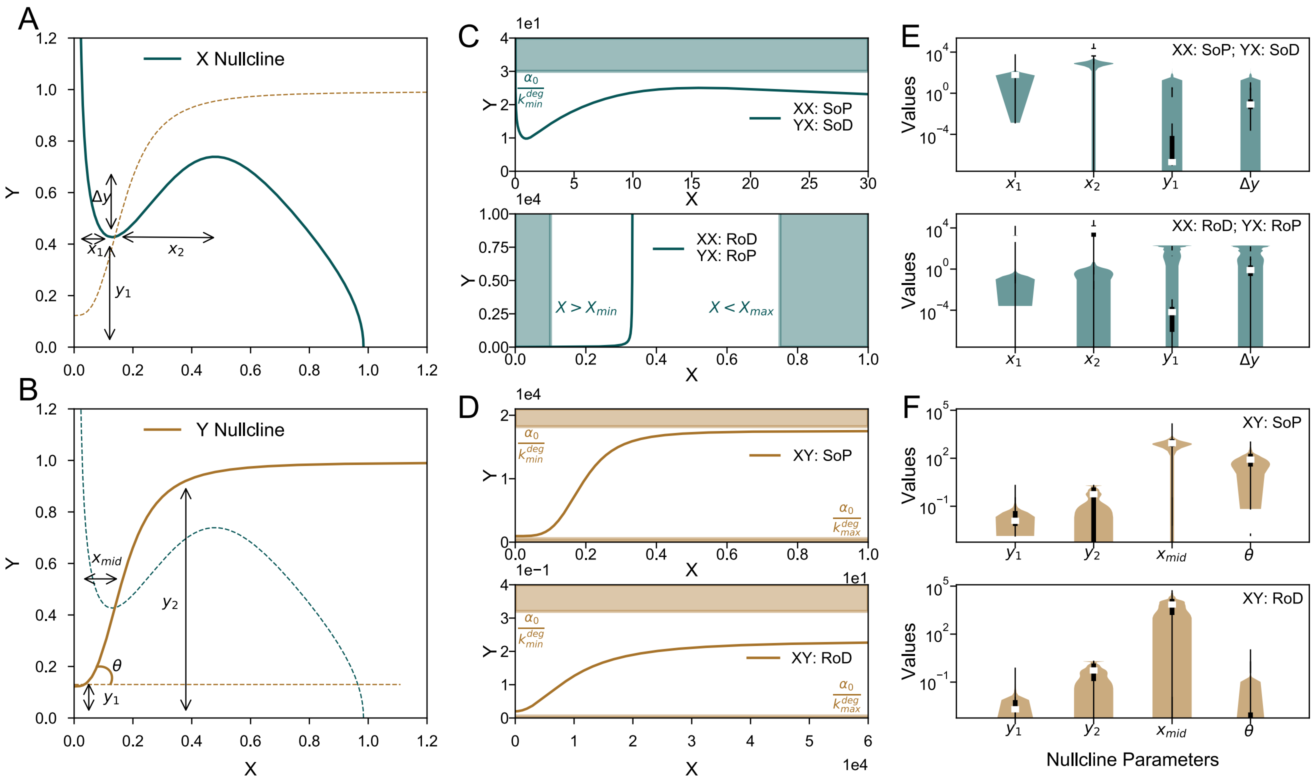


**Fig S2:** To obtain stable limit cycles, the X and Y nullclines must cross obeying the constraints of biochemical reactions. A) Example of an "ideal" X-nullcline geometry that gives stable limit cycles. Limit cycles are obtained when the Y nullcline crosses the X nullcline in the switchback region, defined as the region between the two extrema of the X nullcline. To maximize the likelihood of obtaining limit cycles, the switchback region of the X nullcline, denoted by $x_2$ has to be maximized, $x_1$ has to be minimized, $y_1$ has to be maximized and $\Delta y$ should be minimized. B) shows the different X nullclines obtained by the various XX and YX interactions. The shaded regions correspond to the biochemically forbidden regions of phase-space. C) Represents the distributions of the X nullcline parameters indicated in A), corresponding to the implementations of the interactions at X shown in B i) and B ii). Each panel represents the statistics of the X nullcline parameters over 10,000 random draws. We see that the only implementation that satisfies all the conditions is XX: SoP, YX: SoD. D) Example of an "ideal" Y-nullcline geometry that gives stable limit cycles. To maximize the likelihood of obtaining limit cycles, $x_{mid}$ should be minimized, $y_1$ has to be maximized, and $y_2$ should be minimized, and $\theta$ should be maximized. E) Shows the different Y nullclines obtained by the two XY implementations. F) Each panel represents the statistics of the geometrical quantities of the Y nullcline described in panel B over 10,000 random parameter draws. We see that the only implementation that satisfies all the conditions is XY: SoP.

##### Fig S3. Symmetric and asymmetric systems in GEN Loop Oscillators

To determine the ranges of parameters for numerical simulations, we first simulated the symmetric systems over a wide range of parameters, spanning a few orders of magnitude. For further analyses, we only considered those regions of parameter space, which support stable limit cycles. An example plot of the regions of interest with either $\beta$, or both $\alpha$ and $\beta$ are indicated in color in the top three panels of Fig S3.

We study symmetric and asymmetric systems in the paper. Symmetric systems refers to systems that have the same system parameters for all three dynamical variables, i.e. when $\alpha_1 = \alpha_2 = \alpha_3, \beta_1 = \beta_2 = \beta_3, n_1 = n_2 = n_3, and K_1 = K_2 = K_3$, and so on. Asymmetric systems are as shown in the equations of Appendix Table S3.

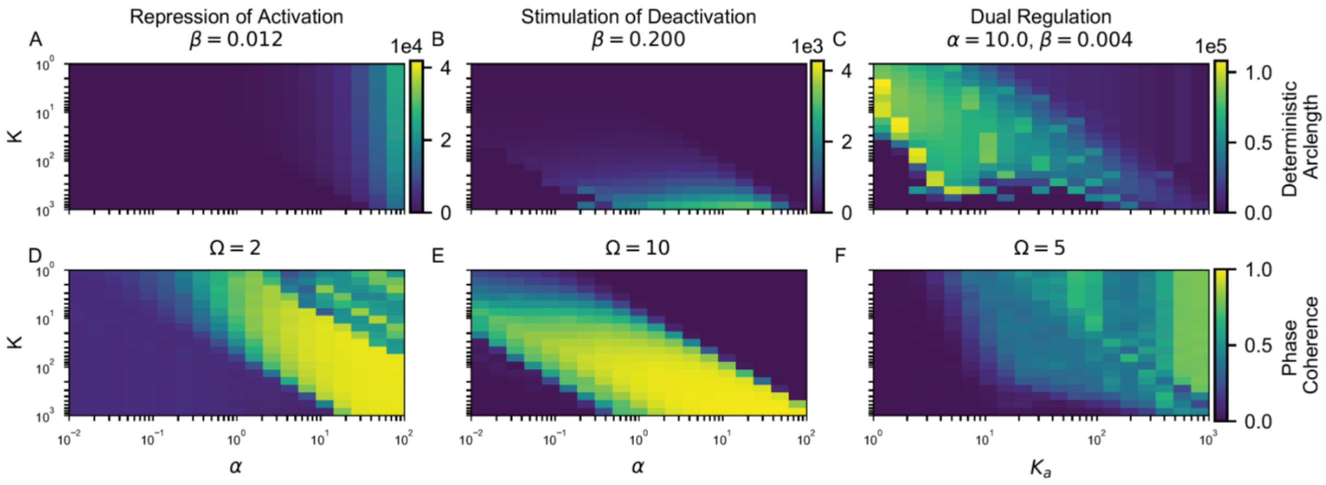


**Fig. S3:** The negative feedback of the repressilator loop is implemented in the three ways discussed in the main text: through repression of Activation (A&D), the Stimulation of Deactivation (B&E), and Dual Regulation (C&F). All three implementations shown are for the symmetric case. In panels A, B, D, E, $\alpha$, and K are varied, keeping $\beta$ fixed. In panels C and F parameters $K_a$,and $K_d$ are varied while keeping $\alpha$ and $\beta$ values fixed. Panels A-C heat plots of the deterministic arclength (color) as the system parameters are varied over a large range. The values we use for further numerical simulations are from the regions that are seen to support stable limit cycle oscillations in the heat plots above. Panels D-F show the phase-coherence of oscillations (color) in the stochastic limits, with system sizes $\Omega$ denoted for each implementation of the repressilator, as system parameters are varied.

#### Fig. S4. Symmetric and asymmetric systems in Post Translational Loop Oscillators

To determine the ranges of parameters for numerical simulations, we first simulated the symmetric systems over a wide range of parameters, spanning a few orders of magnitude. For further analyses, we only considered those regions of parameter space, which support stable limit cycles. These regions are indicated in color in the top three panels of ??.

The typical rates for the production of proteins is experimentally determined as being in the range of a few molecules per min. The distribution we use for the exploration of rates are shown in Table M6. In the plots below, we vary the activation and deactivation rates in each of the three implementations for the PTO loop oscillator shown in ??. The deterministic arclength in each case is indicated by the color in the heatmap shown here.

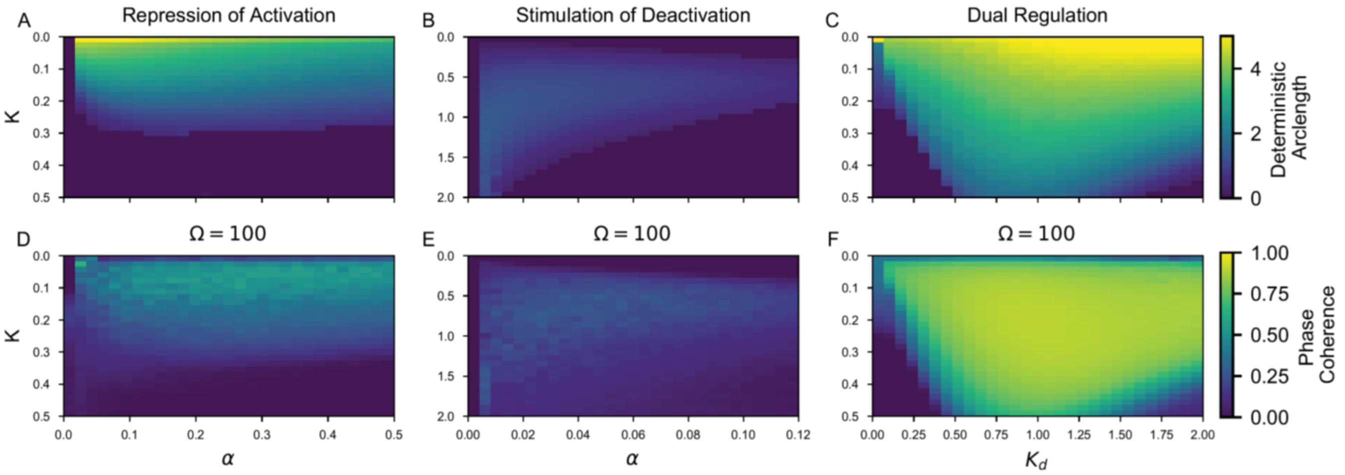


**Fig. S4:** The negative feedback is implemented in three ways in the post translational repressilator loop. Top Panel (L-R): The deterministic arclength is represented as the color for a variation of the two parameters of a symmetric loop oscillator under the (A) Regulation of Activation reaction (B) Regulation of deactivation reaction (C) Dual regulation. All three implementations shown are for the symmetric case. In panels A, B, D, E, $\alpha$, and K are varied, keeping $\beta$ fixed. In panels C and F parameters $K_a$,and $K_d$ are varied while keeping $\alpha$ and $\beta$ values fixed. Bottom Panel (L-R): The phase- coherence calculated numerically through Gillespie SSA is represented as the color for the same variation of parameters as the top panels for the corresponding regulatory implementations.

**Fig S5. Phase-coherence – deterministic arclength relationship in symmetric and asymmetric systems in Post Translational Loop Oscillators**

The negative loop of the repressilator motif, when implemented via dual regulation is the most robust implementation in PTO systems, where the product and degradation are first kinetic order processes.

Implementation of the negative regulation through repressing the activation rate is more robust than implementing the negative regulation by stimulating the deactivation rate. We see that the post translational repressilator motif follows the trends seen in transcriptional repressilator system.

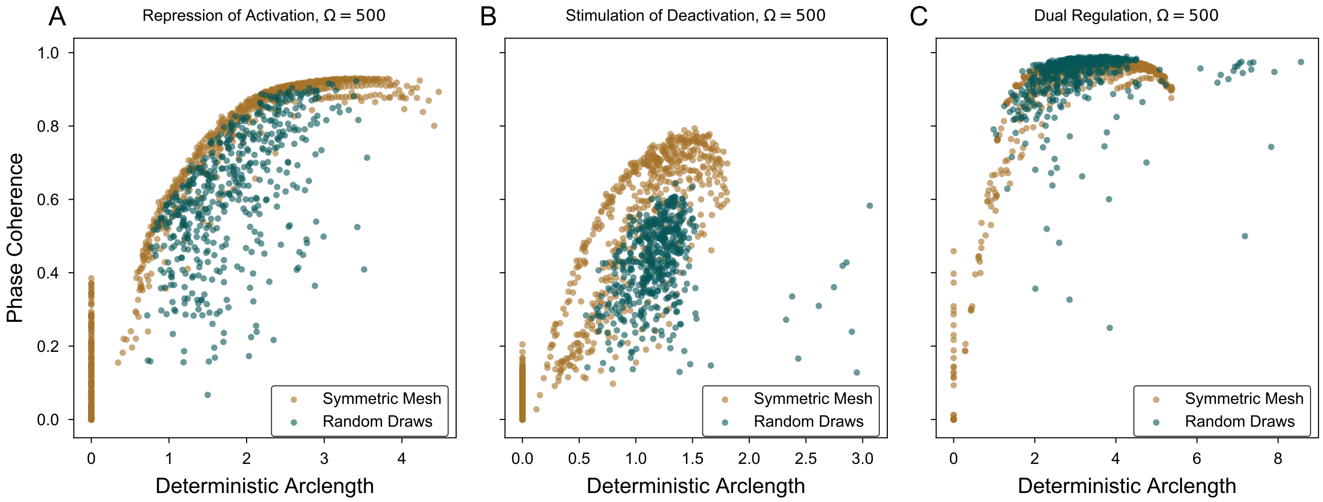


**Fig S5:** Symmetric parameter sets maximize phase-coherence values achieved at a given value of Ω, at a given deterministic arclength, as compared to asymmetric parameter sets in (L-R): (A) Activation regulated, (B) Deactivation regulated, and (C) Dual regulated systems. (D) Fraction of asymmetric parameter sets producing oscillations in the post translational repressilator system.

**Fig S6. Non-dimensionalized Phase Diagram of the loop oscillator to recover the Stimulation of Degradation limit**

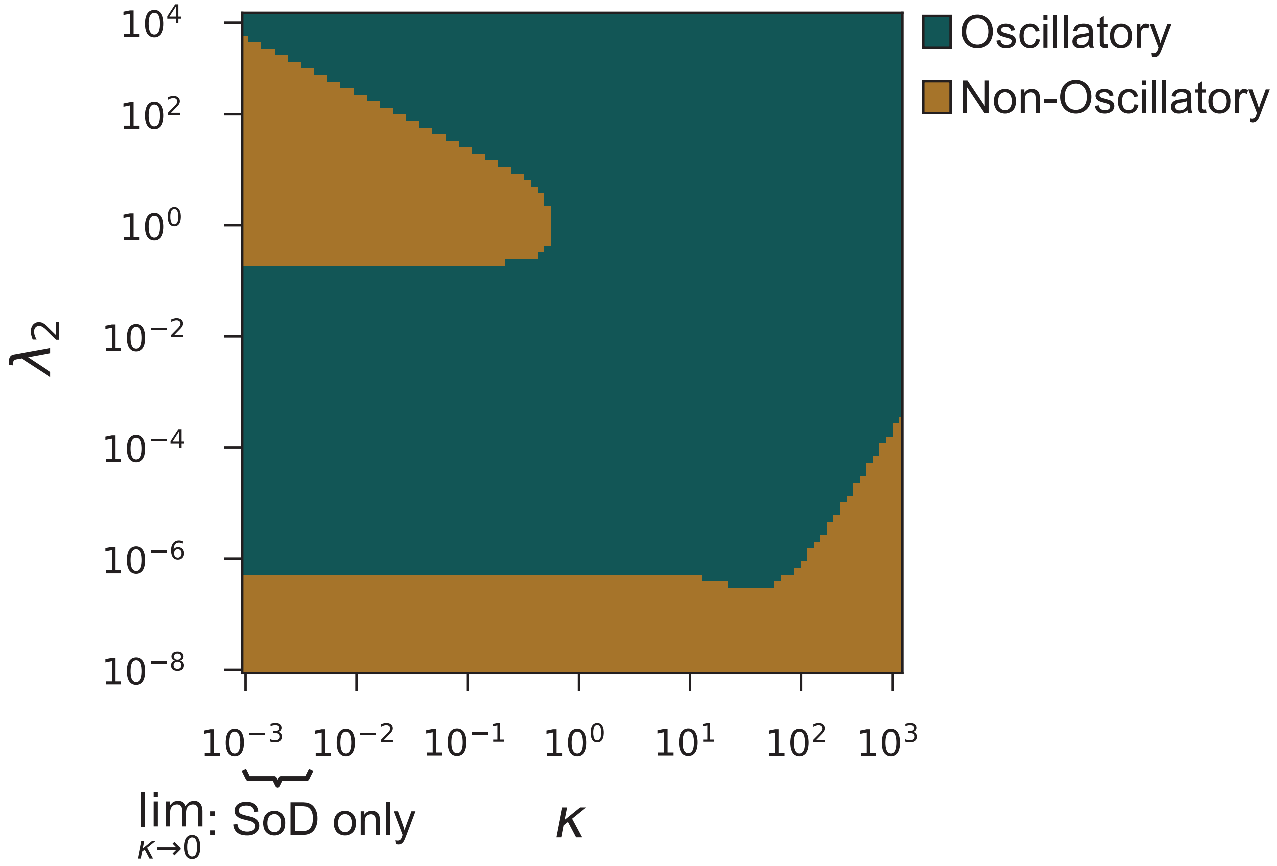


**Fig S6:** Phase diagram depicts the region of stability of a dual regulated repressilator, non-dimensionalized such that the $\kappa \to 0$ limit recovers the regulation through stimulation of degradation mechanism.

Dual Rugulation:

$$\frac{dX}{dt} = \alpha g(Z) - \beta f(Z) X$$
$$\frac{dX}{dt} = \alpha \frac{K_1^n}{K_1^n + Z^n} - \beta \frac{Z^n}{Z^n + K_2^n} X \tag{E.13}$$

When non-dimensionalized such that:

$$x = \frac{X}{K_2}, z = \frac{Z}{K_2}, \kappa = \frac{K_1}{K_2}, \lambda_2 = \frac{}{\beta K_2}, \tag{E.14}$$

We have:

$$\frac{dx}{dt} = \frac{\lambda_2}{1 + ()^n} - \frac{x}{1 + (\frac{1}{z})^n} \tag{E.15}$$

When

$$\lim \kappa \to 0 \frac{dx}{dt} \to \lambda_2 - \frac{z^n}{1 + z^n} x \tag{E.16}$$

we recover the stimulation of degradation only limit of Dual regulation.

**Appendix Table S8. Ranked robustness response of GENs of incoherent input positive-negative oscillators**

| Schematics | # of parameters giving oscillations | Schematics | # of parameters giving oscillations | Schematics | # of parameters giving oscillations |
|---|---|---|---|---|---|
| 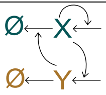 | 4949 | 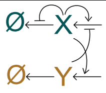 | 31 | 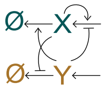 | 0 |
| 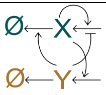 | 889 | 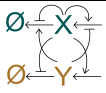 | 19 | 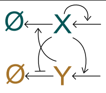 | 0 |
| 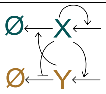 | 343 | 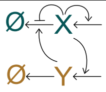 | 13 | 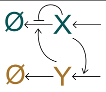 | 0 |
| 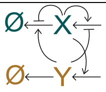 | 287 | 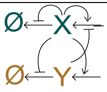 | 11 | 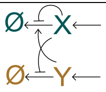 | 0 |
| 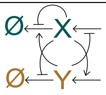 | 128 | 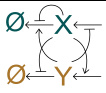 | 10 | 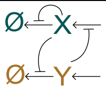 | 0 |
| 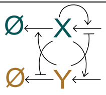 | 127 | 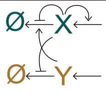 | 1 | 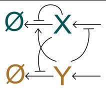 | 0 |
| 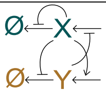 | 66 | 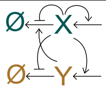 | 1 | 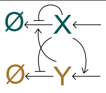 | 0 |
| 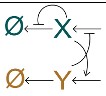 | 55 | 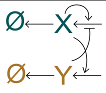 | 0 | 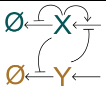 | 0 |
| 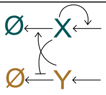 | 37 | 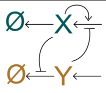 | 0 | 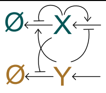 | 0 |

**Appendix Table S8:** The table shows the schematics of the 27 implementations of the incoherent input system, and the corresponding number of parameter sets that support limit cycle oscillations, out of 100,000 parameter sets drawn randomly from the distributions above.

**Appendix Table S9. Ranked robustness response of GENs of coherent input positive-negative oscillators**

| Schematics | # of parameters giving oscillations | Schematics | # of parameters giving oscillations | Schematics | # of parameters giving oscillations |
|---|---|---|---|---|---|
| 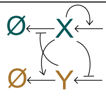 | 4144 | 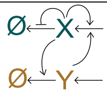 | 57 | 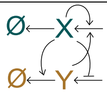 | 0 |
| 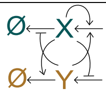 | 539 | 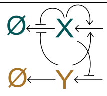 | 42 | 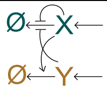 | 0 |
| 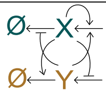 | 504 | 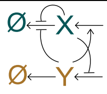 | 38 | 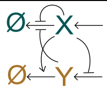 | 0 |
| 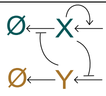 | 315 | 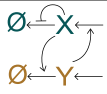 | 10 | 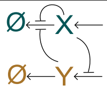 | 0 |
| 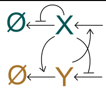 | 213 | 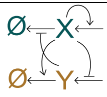 | 8 | 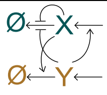 | 0 |
| 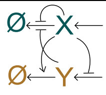 | 110 | 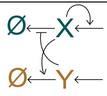 | 6 | 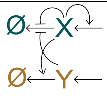 | 0 |
| 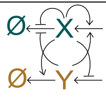 | 103 | 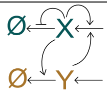 | 2 | 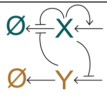 | 0 |
| 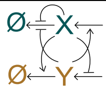 | 97 | 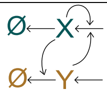 | 0 | 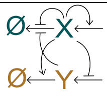 | 0 |
| 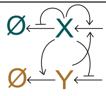 | 83 | 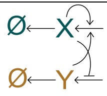 | 0 | 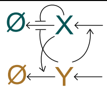 | 0 |

**Appendix Table S9:** The table shows the schematics of the 27 implementations of the coherent input system, and the corresponding number of parameter sets that support limit cycle oscillations, out of 100,000 parameter sets drawn randomly from the distributions above.

**Appendix Table S10. Ranked robustness response of incoherent input post-translational positive-negative oscillators**

| Schematics | # of parameters giving oscillations | Schematics | # of parameters giving oscillations | Schematics | # of parameters giving oscillations |
|---|---|---|---|---|---|
| 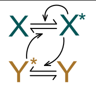 | 234 | 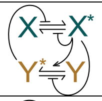 | 3 | 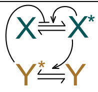 | 0 |
| 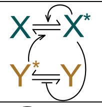 | 38 | 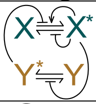 | 2 | 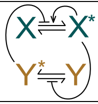 | 0 |
| 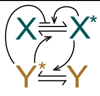 | 25 | 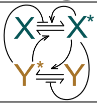 | 1 | 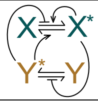 | 0 |
| 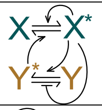 | 20 | 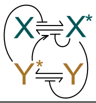 | 1 | 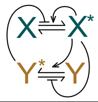 | 0 |
| 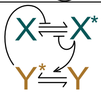 | 14 | 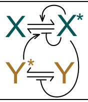 | 1 | 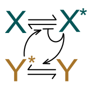 | 0 |
| 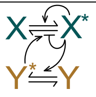 | 8 | 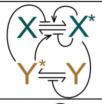 | 1 | 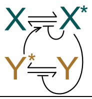 | 0 |
| 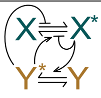 | 5 | 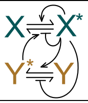 | 1 | 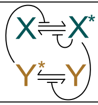 | 0 |
| 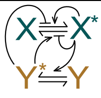 | 5 | 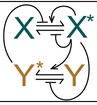 | 1 | 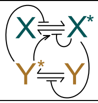 | 0 |
| 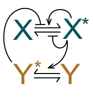 | 4 | 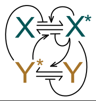 | 1 | 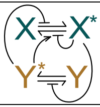 | 0 |

**Appendix Table S10:** The table shows the schematics of the 27 implementations of the post-transcriptional incoherent input system, and the corresponding number of parameter sets that support limit cycle oscillations, out of 100,000 parameter sets drawn randomly from the distributions above.

**Appendix Table S11. Ranked robustness response of coherent input post-translational positive-negative oscillators**

| Schematics | # of parameters giving oscillations | Schematics | # of parameters giving oscillations | Schematics | # of parameters giving oscillations |
|---|---|---|---|---|---|
| 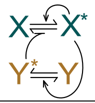 | 91 | 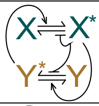 | 2 | 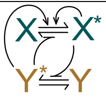 | 0 |
| 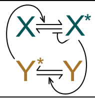 | 64 | 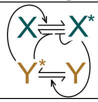 | 2 | 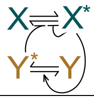 | 0 |
| 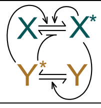 | 44 | 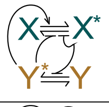 | 1 | 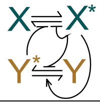 | 0 |
| 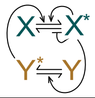 | 10 | 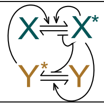 | 1 | 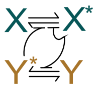 | 0 |
| 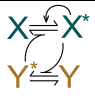 | 5 | 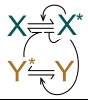 | 1 | 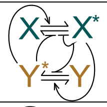 | 0 |
| 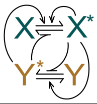 | 5 | 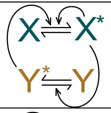 | 0 | 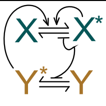 | 0 |
| 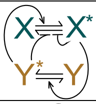 | 5 | 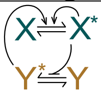 | 0 | 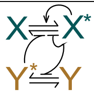 | 0 |
| 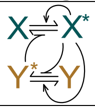 | 4 | 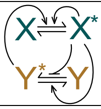 | 0 | 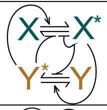 | 0 |
| 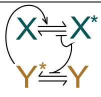 | 4 | 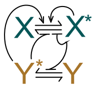 | 0 | 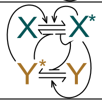 | 0 |

**Appendix Table S11:** The table shows the schematics of the 27 implementations of the post-transcriptional coherent input system, and the corresponding number of parameter sets that support limit cycle oscillations, out of 100,000 parameter sets drawn randomly from the distributions above.
